# Multiplexed volumetric CLEM enabled by antibody derivatives provides new insights into the cytology of the mouse cerebellar cortex

**DOI:** 10.1101/2023.05.20.540091

**Authors:** Xiaomeng Han, Xiaotang Lu, Peter H. Li, Shuohong Wang, Richard Schalek, Yaron Meirovitch, Zudi Lin, Jason Adhinarta, Daniel Berger, Yuelong Wu, Tao Fang, Elif Sevde Meral, Shadnan Asraf, Hidde Ploegh, Hanspeter Pfister, Donglai Wei, Viren Jain, James S. Trimmer, Jeff W. Lichtman

## Abstract

Mapping neuronal networks that underlie behavior has become a central focus in neuroscience. While serial section electron microscopy (ssEM) can reveal the fine structure of neuronal networks (connectomics), it does not provide the molecular information that helps identify cell types or their functional properties. Volumetric correlated light and electron microscopy (vCLEM) combines ssEM and volumetric fluorescence microscopy to incorporate molecular labeling into ssEM datasets. We developed an approach that uses small fluorescent single-chain variable fragment (scFv) immuno-probes to perform multiplexed detergent-free immuno-labeling and ssEM on the same samples. We generated eight such fluorescent scFvs that targeted useful markers for brain studies (green fluorescent protein, glial fibrillary acidic protein, calbindin, parvalbumin, voltage-gated potassium channel subfamily A member 2, vesicular glutamate transporter 1, postsynaptic density protein 95, and neuropeptide Y). To test the vCLEM approach, six different fluorescent probes were imaged in a sample of the cortex of a cerebellar lobule (Crus 1), using confocal microscopy with spectral unmixing, followed by ssEM imaging of the same sample. The results show excellent ultrastructure with superimposition of the multiple fluorescence channels. Using this approach we could document a poorly described cell type in the cerebellum, two types of mossy fiber terminals, and the subcellular localization of one type of ion channel. Because scFvs can be derived from existing monoclonal antibodies, hundreds of such probes can be generated to enable molecular overlays for connectomic studies.

## Introduction

Mapping neuronal networks that underlie behavior is a central focus in neuroscience. Techniques that reveal the structure and molecular components of neurons and their connections have expanded over the last several decades. Automated high resolution serial section electron microscopy (ssEM) provides detailed structural information about neuronal networks but omits other important features of neuronal ensembles. For example, the molecular markers of cell or synapse types, the localization of specific molecules related to neuronal physiology, or the functional properties of neural circuits cannot be provided by ssEM alone. Therefore, knowledge gained from neuronal networks mapped by ssEM is still limited. There is a need for new technology to incorporate molecular and functional information into ssEM.

Several techniques that are useful for obtaining both structural context and molecular information in the same sample use light and electron microscopy. For light microscopy (LM), methods to colorize cells, such as Brainbow, take advantage of combinatorically expressed fluorescent proteins to enable dense labeling of neurons with multiple, distinct colors ^1, 2^. However, the resolution of fluorescence imaging is limited by diffraction. This limitation makes it challenging to differentiate among densely packed neurites or to assign the pre- and postsynaptic partners of each synapse. Expansion microscopy (ExM) can overcome this limitation ^3^ by physically expanding the dimensions of a sample. When combined with fluorescent lipid labeling ^4^, ExM alone or combined with super resolution approaches, have the potential to yield both structural and functional information in the same neural sample ^5, 6^. Whether these all-fluorescence approaches will ultimately have comparable speed and resolution as electron microscopy is still uncertain. At present nanoscale electron microscopy is the gold standard for imaging cellular structure and any approaches that combine it with multi-molecular labeling would add greater interpretive power to this tried-and-true technique. To label specific molecules directly in electron microscopy images antibodies can be applied before either before or after resin embedding. The antibodies are visible by virtue of being tagged with metal beads or enzymes that chemically create electron-dense deposits visible by EM ^7–9^. Antibodies conjugated to metal beads of different sizes allow identification of up to three different molecular labels ^9^. However, these immuno-labeling strategies have several technical challenges. Pre-resin embedding immunolabeling requires membrane permeabilization for antibody access that perforates cell membranes, causing them to appear discontinuous in EM images. Discontinuous membranes are incompatible with neural circuit tracing. Post-embedding methods (typically after sectioning) suffer from both the loss of antibody binding due to denatured antigen epitopes by the heavy metal staining and the challenges of using aqueous immunoreagents with hydrophobic resin embedded sections.

One way to circumvent the access problem associated with pre- or post-embedding is to avoid antibodies altogether. Molecular labeling via transgenically engineered probes with EM-visible molecular tags like miniSOG ^10^ or APEX ^11^ can be used to label one or more cell-types with ssEM ^12, 13^, the current maximum number of labels being four ^13^. This technical limitation in the number of different labels that can be combined in transgenic approaches suggests that more scalable methods would be of value.

One such approach is volumetric correlated light and electron microscopy (vCLEM) in which fluorescence LM and ssEM are performed sequentially on the same sample and subsequently co-registered. Because multiplex labeling (e.g. with many colors) can easily be achieved by fluorescence LM sing spectral unmixing ^14, 15^, vCLEM can potentially incorporate more molecular and functional information into ssEM datasets than by directly tagging within the electron microscopy images. Moreover, vCLEM performed on animals that express a fluorescent protein or have a calcium indicator can identify certain cell types ^16, 17^ or show the connectivity patterns of functionally identified cells ^18–22^. The challenge of vCLEM with using conventional antibodies for immunofluorescence is, as mentioned above, the requirement of permeabilizing detergents for the antibodies to gain access to intracellular sites. These detergents compromise membrane structure ^23^. Attempts to circumvent the need for permeabilization include the post-resin embedding techniques used in array tomography in which immunolabeling of partly hydrophilic resin serial sections is followed by modified heavy metal staining (to maintain antigenicity of the tissue) and EM imaging ^24^. This CLEM approach is effective, but the modified heavy metal staining compromises ultrastructure and the water-permeant resin is too soft for the ultrathin (30-40nm) sectioning needed for connectomics.

Detergent-based permeabilization is also avoided by use of nanobodies as immunolabels ^23^. Nanobodies are single domain immuno-binders derived from camelid heavy chain only antibodies ^25^. Because nanobodies are 1/10 the size of conventional antibodies, they diffuse into tissue samples without permeabilizing agents like Triton X-100. In the presence of preserved extracellular space (ECS), nanobodies diffused over a distance of ∼100 micrometers into brain samples within 48 hours of their application ^23^. Nanobodies, however, for brain markers are uncommon and immunizing camelids is time-consuming to do. ECS preservation also facilitates standard antibody (IgG) diffusion into brain tissue samples without detergent treatment ^26, 27^. However, fluorescent labeling with IgGs is challenging if one desires multiple color probes in the same sample. The traditional approach of using fluorescent secondary antibodies to amplify the signal requires that each primary antibody originates from a different species. This limits the number of standard antibodies that can be disambiguated in one tissue sample. Alternatively, the full-length antibody can be tagged directly with fluorophores that are covalently linked to a particular amino acid moiety. However, given the likelihood of many binding sites this approach can hinder the avidity of the probe and cause different antibodies to the same epitope to have different intensities obviating quantitative assays of fluorescence intensity. The goal of this study is to develop a larger number of immuno-probes that diffuse across cell membranes in the absence of detergents and combine them with multiplex imaging techniques like spectral unmixing for both cell wide and subcellular localization of epitopes.

Our CLEM approach is to generate small immuno-probes from full-size IgG monoclonal antibodies. Monoclonal antibodies (mAbs) are homogeneous with respect to their amino acid sequence and the epitope recognized ^28^. Creating smaller probes from existing mAbs requires knowledge of a particular mAb’s amino acid sequence. These sequences can be obtained by cDNA cloning from the hybridoma that produces the mAb ^29–32^. Alternatively, if the hybridoma cells are not available, the amino acid sequence can be derived from the mAb protein itself ^33, 34^. With the knowledge of the amino acid sequence of a mAb, it is possible to synthesize a smaller single-chain variable fragment (scFv) that retains the binding specificity of the progenitor mAb. ScFvs are built by recombinantly linking the V_H_ and V_L_ domains of mAbs via a flexible peptide linker (Figure 1 a) ^35^. Only if the V_H_ and V_L_ pair correctly (∼60% of the time) do they bind to the antigen ^30, 35^. Because a scFv consists of only V_H_ and V_L_ domains, they are 1/5 the size of conventional antibodies. We surmised that their small size would allow them to diffuse into tissue samples without detergent-based permeabilization, as is the case for nanobodies ^23^. Because there exist extensive collections of well-characterized mAbs that selectively label neuronal cell types or signaling molecules ^36^, it is possible to develop a large collection of the corresponding scFvs for use in the nervous system. Because scFvs are produced recombinantly ^30^, they can be engineered so that different fluorescent dyes can easily be conjugated to them for multiplex imaging.

**Figure 1.**
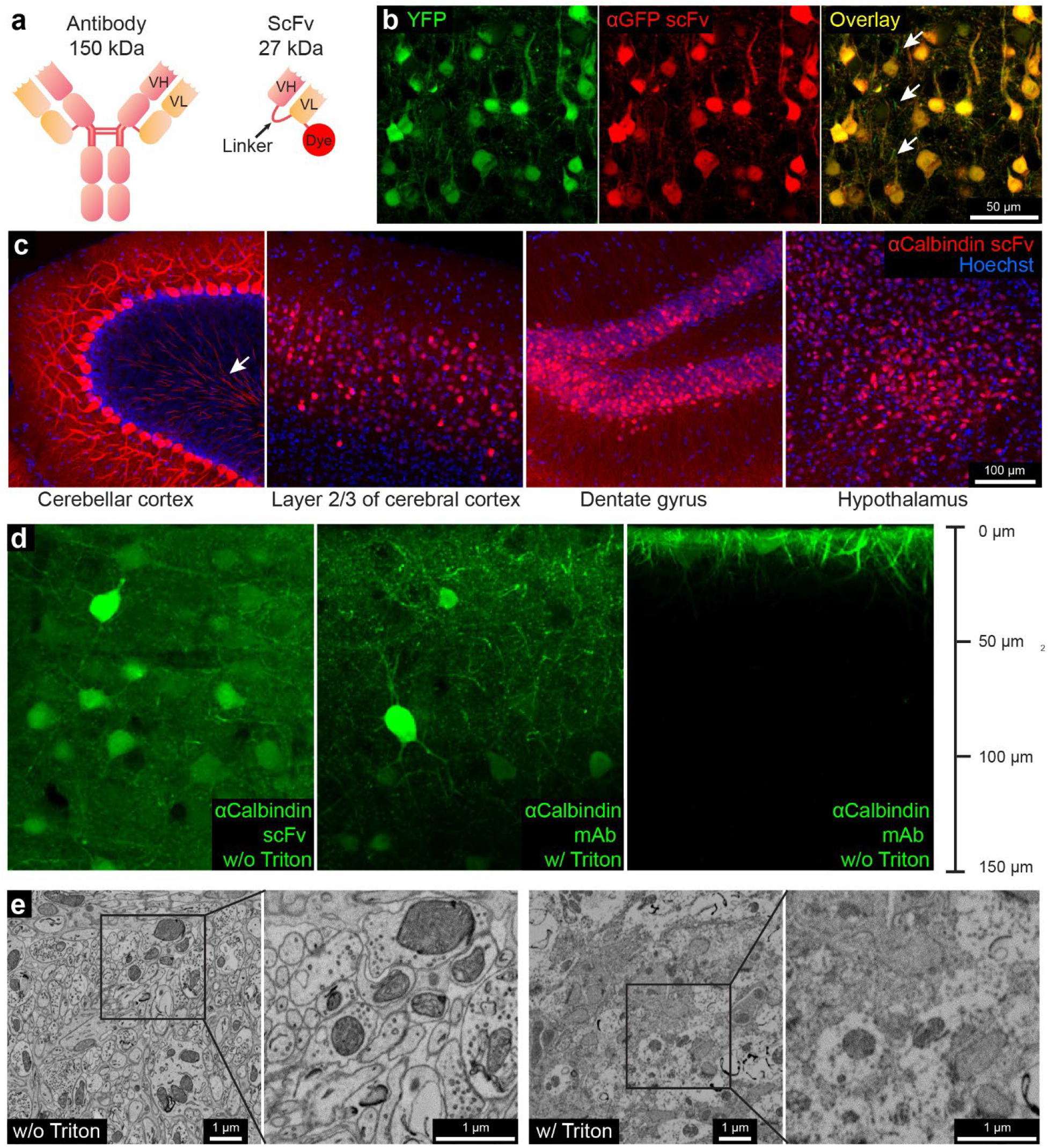
Fluorescent scFv probes label brain tissues without detergents to preserve electron microscopy ultrastructure. **a**, Schematic representations of a full-length IgG antibody and an scFv probe with a conjugated fluorescent dye. **b**, Confocal images from the cerebral cortex of a YFP-H mouse labeled using a GFP-specific scFv probe conjugated with the red dye 5-TAMRA. Arrows show thinner neuronal processes, perhaps myelinated, that are not labeled by scFv. **c**, Brain regions labeled with a calbindin-specific scFv probe conjugated with 5-TAMRA. The arrow in the left panel shows myelinated Purkinje cell axons in the granule layer. **d**, Tissue penetration depth comparison of scFvs, mAbs, and the role of detergents in improving the depth of labeling. **e**, Comparison of ultrastructure with and without Triton X-100 on scFv labeled samples. Boxed insets are shown at higher magnification in adjacent panels.

To test this idea, we developed eight scFvs based on eight well-characterized mAbs and conjugated them with various fluorescent dyes (see Table 1). Each scFv in this panel proved effective as a detergent-free immunofluorescent probe. We then used linear unmixing with confocal microscopy to visualize six different functional molecular markers in the same brain sample and co-registered these labels to ssEM volumes with pristine preservation of the ultrastructure of the same samples. Because the volumetric fluorescent and electron microscopy image data were of excellent quality, we believe this approach holds promise for routine linking of molecular information to connectomic information obtained from the same tissue samples.

**Table 1.** Fluorescent scFv immuno-probes generated in this work.

| **Target** | **mAb Clone No.** | **Fluorescent dyes**<br>**conjugated** | **Link of expression plasmid at**<br>**Addgene** |
| --- | --- | --- | --- |
| **GFP** | N86/38 | 5-TAMRA | https://www.addgene.org/204419/ |
| **Calbindin** | L109/57 | 5-TAMRA, Alexa Fluor<br>488 | https://www.addgene.org/204421/ |
| **GFAP** | N206B/9 | 5-TAMRA | https://www.addgene.org/204420/ |
| **VGluT1** | N28/9 | 5-TAMRA, Alexa Fluor<br>532 | https://www.addgene.org/204424/ |
| **PSD-95** | K28/43 | 5-TAMRA | https://www.addgene.org/204422/ |
| **Kv1.2** | K14/16 | Alexa Fluor 594 | https://www.addgene.org/204425/ |
| **Parvalbumin** | L114/81 | Alexa Fluor 647 | https://www.addgene.org/204423/ |
| **Neuropeptide Y** | L115/13 | Alexa Fluor 647 | https://www.addgene.org/204426/ |

## Results

### Generation of scFv-based immuno-probes and their use in detergent-free immunofluorescence

To test the hypothesis that scFvs could work as immuno-probes for detergent-free immunofluorescence, we first generated an anti-green fluorescent protein (GFP) scFv based on the amino acid sequence of the anti-GFP mouse mAb N86/38 ^36^. This mAb binds to both GFP and the GFP derivative YFP. The sequence (derived by PCR cloning from hybridoma cells ^29^ was used to construct an scFv in which the V_H_ and the V_L_ domains were connected via a 3x linker (GGGGS GGGGS GGGGS) (Figure 1 a). A sortase tag ^23^ was added to the C-terminus of the V_L_ domain of the scFv for dye conjugation, followed by a 6 x His tag for purification. After the scFv was expressed in Expi 293 cells and purified by affinity chromatography, the fluorescent dye 5-TAMRA was linked (see Methods) (Figure 1 a). We tested this red fluorescent anti-GFP scFv probe with our detergent-free immunofluorescence protocol (see Methods) on cerebral cortex from YFP-H mice ^37^ (n=3). The images showed colocalization of fluorescence signals from native YFP and red fluorescence from the scFv probe (Figure 1 b; Sup. Figure 1 a). The scFv probe thus retains the binding properties of the parental mAb and penetrates aldehyde-fixed brain tissue without the need for detergent treatment. Although the fluorescent signals colocalized in cell bodies and thick dendrites, a few thinner neuronal processes, possibly myelinated axons, were not well labeled with the scFv (arrows in Figure 1 b; but see below).

To study the depth of penetration, we immunolabeled two 300-μm thick samples from mouse cerebral cortex without detergent treatment (see Methods). After a free-floating incubation of seven days, this scFv penetrated to a depth of ∼60 μm into the tissue (Sup. Figure 1 b). This was deeper than the 10-μm penetration of a fluorescently labeled anti-GFP polyclonal antibody (pAb) (Sup. Figure 1 b). Furthermore, we could improve the penetration to >100 μm by use of an ECS preserving perfusion protocol ^27^ (Sup. Figure 1 c). Fluorescently labeled nanobodies, which are half the size of scFvs, penetrate even deeper in the same time (Sup. Figure 1 b, c).

We then proceeded to produce an scFv probe for an endogenously expressed protein (calbindin, also known at calbindin-D 28k). This probe was based on a well-characterized mouse mAb (L109/57; ^36^). Calbindin (CB) is a calcium-binding protein expressed by certain neuronal types in various brain regions, such as cerebellar Purkinje cells, neurons in layer 2/3 of the cerebral cortex, dentate gyrus granule cells, as well as a subpopulation of hypothalamic neurons ^38^. Because previous studies showed that an scFv’s performance improved as the length of linker increased ^39, 40^, we chose a 4x linker (GGGGS GGGGS GGGGS GGGS) rather than a 3x linker. We conjugated the anti-CB scFv to fluorescent dyes and tested it on brain tissue samples (n=2) from wild type C57BL/6J mice with the detergent-free immunofluorescence protocol. This scFv probe labeled neurons in all four brain regions where CB-expressing neurons were highly abundant (Figure 1 c) showing that these probes can label endogenous protein targets via detergent-free immunofluorescence. The anti-CB scFv labeled many neuronal processes in the cerebellar granule layer (arrow in Figure 1 c, first panel). Presumably these processes are the myelinated axons of the labeled Purkinje cells. The stronger labeling of cellular processes obtained with this scFv when compared to the anti-GFP scFv, may be explained by the different lengths of the linkers used. We repeated the penetration test (n=2) (see Methods) for this probe and found that the anti-calbindin scFv penetrated to a depth of ∼150 μm in a 300-μm tissue slice, a distance beyond the maximum imaging depth of confocal microscopy without tissue clearing (Figure 1 d, first panel). This is the same penetration depth as seen for the L109/57 mAb from which the scFv is derived when using a conventional immunofluorescence protocol ^41^, with Triton X-100 as the detergent for permeabilization (Figure 1 d, the first and second panels). In stark contrast, the mAb only penetrates the surface of a tissue sample prepared without detergent (Figure 1 d, the third panel). The improved binding afforded by the use of a longer linker may have given rise to observable signals deeper in the specimen when compared to the anti-GFP scFv.

To evaluate the ultrastructure in the anti-CB scFv labeled cerebellum, we performed a routine EM staining protocol (see Methods) on a 120-μm thick cerebellar cortex sample that had been treated for 7-days with the free-floating immunofluorescence protocol both with and without the use of Triton X-100 for membrane permeabilization (see Methods). The ultrastructure of the sample without Triton X-100 was well-preserved, with continuous membranes and clearly visible synaptic vesicles (Figure 1 e, the first and second panels). The samples with Triton X-100, in contrast, generated EM images in which membrane structures were discontinuous and synaptic vesicles no longer identifiable (Figure 1 e, the third and fourth panels).

### Generation of fluorescent scFv probes targeting various targets

To achieve multiplex labeling with scFvs, we included six additional scFvs (anti-vesicular glutamate transporter 1 (VGluT1), anti-glial fibrillary acidic protein (GFAP), anti-voltage-gated potassium channel subfamily A member 2 (Kv1.2), anti-parvalbumin (PV), anti-postsynaptic density protein 95 (PSD-95), and anti-neuropeptide Y (NPY)). These scFvs are based on six well-characterized mouse mAbs ^36^ (see Table 1). We conjugated them with various fluorescent dyes via the sortase reaction to create fluorescent immuno-probes (see Table 1). We then validated them by comparing scFv versus parental mAb immunofluorescence for each probe (Sup. Figures 2-4). Probes for CB, VGluT1, GFAP, Kv1.2, and PV generated detergent-free immunofluorescence patterns that were similar to or in some cases stronger than their parental mAbs in Crus 1 of the cerebellar cortex (Figure 2 a; Sup. Figure 2; Sup. Figure 3). For example, the scFvs that target CB and PV produced more consistent labeling results for the cell bodies and nuclei of Purkinje cells than labeling with full-size mAbs, which never labeled the nuclei of Purkinje cells (Sup. Figure 2 a, e). Nonetheless, nuclear labeling varied between cells (Sup. Figure 2 a, e). This variability may be attributable to variable expression of CB and PV in the target cells, as previously shown for Purkinje cells in different lobules ^42, 43^. Levels of CB and PV are variable in both the cell nucleus and cytoplasm of Purkinje cells ^44, 45^. The axons of Purkinje cells were well labeled with both anti-CB and anti-PV scFvs (Sup. Figure 2 a, e; Sup. Figure 3), unlike low levels of axon labeling with these full-size antibodies. This difference may be a consequence of impaired access of full-size antibodies to myelinated axons. Post-embedding labeling and thin cryosections have shown that both PV and CB are expressed in axons ^46, 47^, confirming the results shown here with scFvs. Probes for PSD-95 and neuropeptide Y (NPY) showed results similar to those found with full size mAbs (Sup. Figure 4), but only when labeling was done on tissue samples where the ECS was preserved.

**Figure 2.**
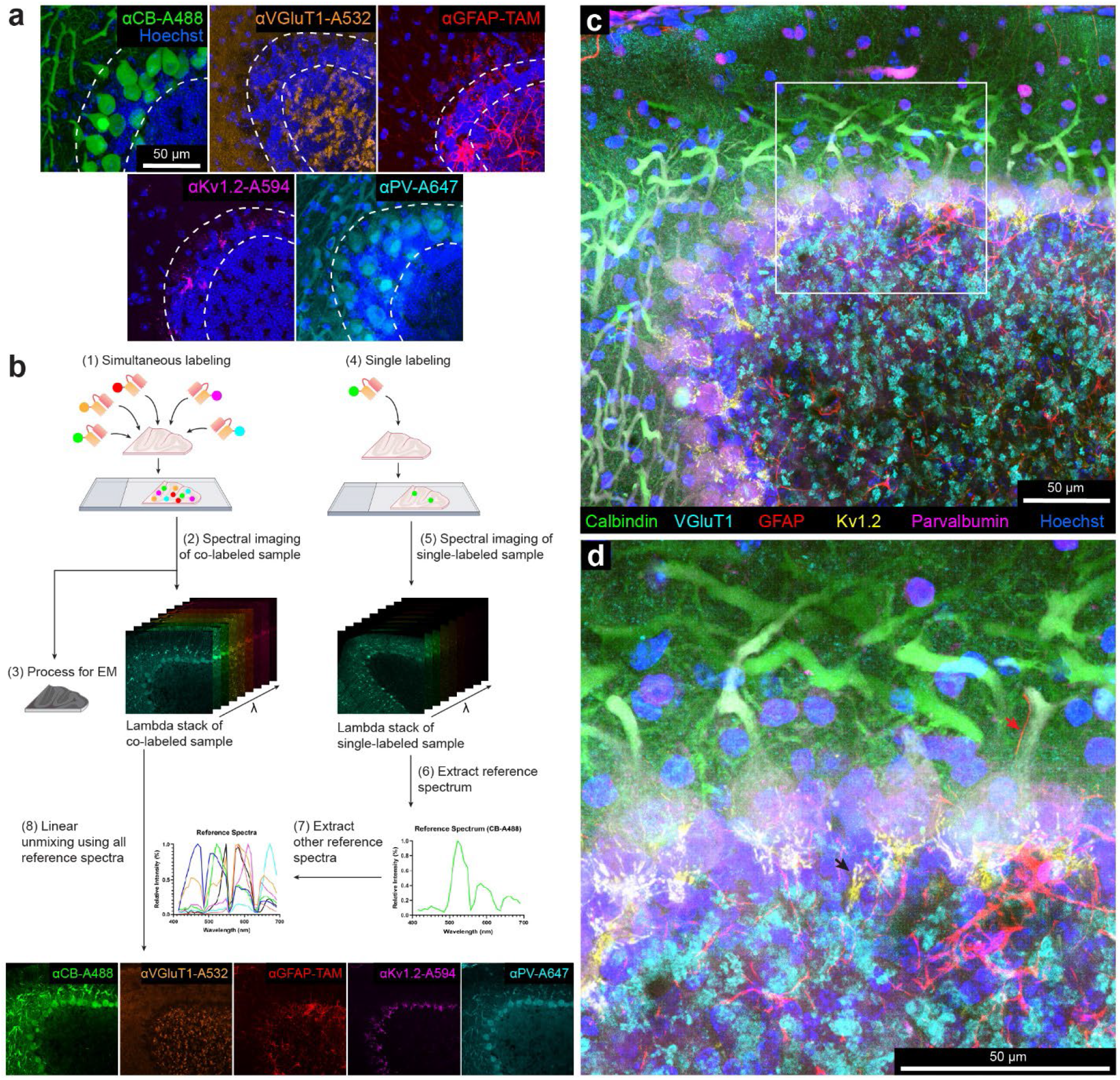
Multi-color immunofluorescence enabled by scFv probes and linear unmixing of confocal micrographs. **a**, Confocal images of different sections from the cerebellum labeled with: a calbindin-specific scFv probe conjugated with Alexa Fluor 488, a VGluT1-specific scFv probe conjugated with Alexa Fluor 532, a GFAP-specific scFv probe conjugated with 5-TAMRA, a Kv1.2-specific scFv probe conjugated with Alexa Fluor 594, and a parvalbumin-specific scFv probe conjugated with Alexa Fluor 647. The double dotted lines delineate the Purkinje cell layer. (see Sup. Figure 2 for larger fields of view). CB, calbindin; VGluT1, vesicular glutamate transporter 1; GFAP, glial fibrillary acidic protein; Kv1.2, voltage-gated potassium channel subfamily A member 2; PV, parvalbumin; TAM, 5-TAMRA. **b**, Workflow of multi-color imaging enabled by scFv probes and linear unmixing (see text). **c**, Maximum intensity projection of the multi-color fluorescence image stack acquired by linear unmixing of confocal images. The signal of each fluorescent dye was pseudo-colored for better visualization. **d**, Enlarged boxed inset from **c**. Red arrow indicates a Bergmann fiber (GFAP-positive) adjacent to the main dendrite of a Purkinje cell. Black arrow indicates sites where axons form a pinceau structure labeled by the Kv1.2-specific scFv probe.

### Multiplexed vCLEM based on scFv-enabled immunofluorescence using linear unmixing

To attempt multiplexed vCLEM, we chose five fluorescent probes with different targets and linked to different fluorophores (anti-CB-Alexa Fluor 488, anti-VGluT1-Alexa Fluor 532, anti-GFAP-5-TAMRA, anti-Kv1.2-Alexa Fluor 594, and anti-PV-Alexa Fluor 647) to simultaneously label a 120-μm thick cerebellar sample from Crus 1 (Figure 2 b, see Methods). Because these fluorophores have overlapping excitation and emission spectra (Sup. Figure 5), we used spectral unmixing on a laser-scanning confocal microscope for multiplex imaging. A lambda stack of the multicolor sample with a depth of 52 μm was imaged by confocal microscopy (Zeiss LSM 880) using a spectral detector (Figure 2 b; Sup. Figure 5 b). Because a fluorophore’s reference spectrum (required for linear unmixing) may vary depending on brain regions ^15^ we acquired a lambda stack for each fluorophore in individually labeled samples from the same region as the multi-labeled sample (Figure 2 b; Sup. Figure 5 c). Using a linear unmixing algorithm and the reference spectra of all five dyes (Figure 2 b; Sup. Figure 5 c), the lambda stack of the multi-labeled sample was separated into five fluorescent probe channels (Figure 2 b; Sup. Figure 6 a). In some cases, spectral unmixing can be accomplished more efficiently by extracting reference spectra from the multi-labeled sample ^48^, but these approaches were unsatisfactory with this sample (Sup. Figure 6 b, c).

The spectrally unmixed labeling pattern for each probe was distinct and resembled what we found in the individually labeled samples (Figure 2 c; Sup. Figure 6 a; Sup. Figure 7). In our sample cell nuclei stained with Hoechst dye) was acquired without spectral unmixing using 405-nm laser excitation. The short excitation and emission wavelength of Hoechst scattered strongly and lost intensity dramatically with depth requiring a 10-fold increase in laser power for the deepest parts of the volume but we found in a separate identically prepared sample, Hoechst could be combined with the other linear unmixing channels with equally good results (Sup. Figure 8). Overall, we used linear unmixing to acquire a multi-color fluorescence image volume (304 μm x 304 μm x 52 μm), showing the labeling of the five scFv probes (Figure 2 c, d; Sup. Figure 9). Each fluorescence channel is highly specific for the labeling pattern of the corresponding scFv. For example, the labeling of Bergmann fibers (red arrow) by the anti-GFAP scFv and the labeling of the axons in pinceau structures (black arrows) by the anti-Kv1.2 scFv are easily distinguishable (Figure 2 d). After confocal imaging was completed, the sample was stained with heavy metals and embedded in resin (see Methods) and an isotropic resolution (voxel size 1.15 μm^3^) image volume of the sample was acquired by μCT (see Methods).

The sample was then cut into ∼4000 serial 30 nm ultrathin sections using an automated tape-collecting ultramicrotome ^49^. A low-resolution overview image subvolume of 1 mm x 1 mm x 60 μm with a voxel size of 150 nm x 150 nm x 1.35 μm, was obtained with a single beam scanning electron microscope from the superficial half of all the serial sections where confocal imaging was of high quality. This was followed by multi-beam scanning electron microscope imaging to collect a high-resolution volume of 164 μm x 200 μm x 26 μm at a resolution of 4 nm x 4 nm x 30 nm (Figure 3 a), comprising the superficial 22% of the serial sections. Tissue ultrastructure appeared well preserved throughout the 60-μm subvolume that included all layers of the cerebellar cortex (Figure 3 a, 1-4; Sup. Figure 10).

**Figure 3.**
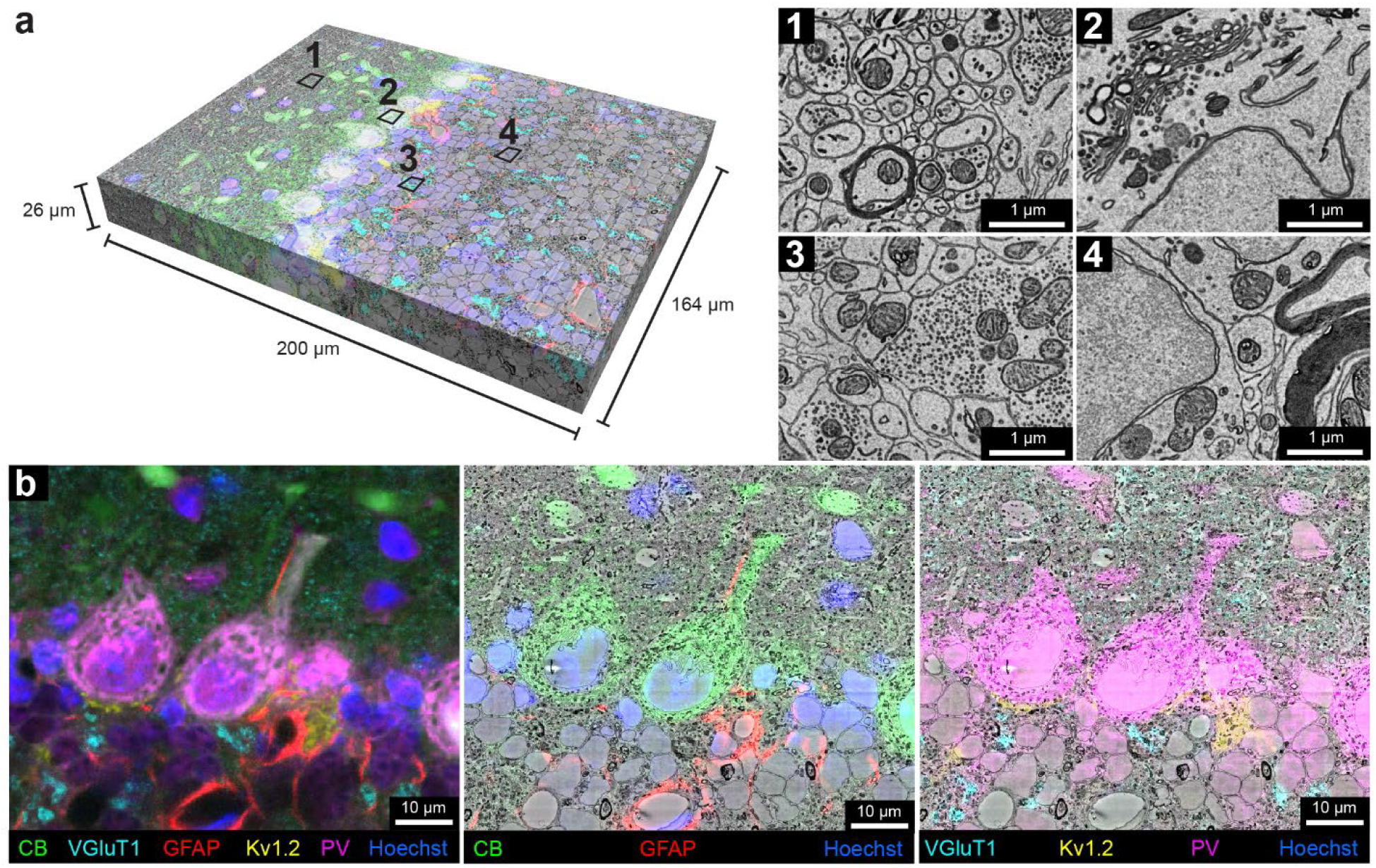
Multi-color volumetric CLEM enabled by scFv-assisted immunofluorescence. **a**, The high-resolution ssEM volume acquired from the cerebellar lobule, Crus 1 with multi-color immunofluorescence from scFv probes separated by linear unmixing. The multi-color fluorescence data was co-registered with the high-resolution ssEM data. Numbers **1**-**4** indicate approximate regions where the ultrastructure was examined at high resolution. Owing to the absence of detergent in immunofluorescence labeling, fine ultrastructure was preserved throughout the ssEM volume, such as in the molecular layer (**1**), in the Purkinje cell layer (**2**), in the glomeruli in the granule cell layer (**3**), and in the granule cell bodies (**4**). **b**, Demonstration of the overlay between fluorescence signals and EM ultrastructure. Left panel shows the multi-color six-channel fluorescent image of slice 250 (of 848) of the spatially transformed fluorescence image volume. Middle panel shows three fluorescence channels corresponding to the labeling of CB, GFAP, and Hoechst overlaid onto the EM micrograph of slice 250. Right panel shows four fluorescent channels corresponding to the labeling of VGluT1, Kv1.2, PV, and Hoechst overlaid onto the EM micrograph of slice 250.

Co-registration between the stacks of fluorescence images and EM volumes was accomplished by visually finding common fiduciaries (such as blood vessels) in the μCT volume and confocal volume as an intermediate step to guide the trimming of the block to the relevant region prior to ultra-thin sectioning (see Methods). This morphing registration overcame the rotation, tilt, stretch, and warping that distort the electron microscopy sections ^50^. To target the location for high-resolution EM imaging, the confocal stack to the low-resolution EM stack were registered. For the final, high-precision co-registration between the six-color fluorescence confocal volume and the high-resolution EM volume, 188 manually picked landmark points that corresponded to the same sites in the two volumes were placed on blood vessels, cell nuclei, cell bodies, and axons (Sup. Figure 11). A 3D transformation of the fluorescence volume was produced by a thin plate spline interpolation based on the point correspondences (BigWarp in FIJI) ^51^. The transformed fluorescence volume was then overlaid with the high-resolution EM volume to create the vCLEM dataset (Figure 3 a). The fluorescence signals from the molecular markers corresponded with objects visible in the EM ultrastructure throughout the volume (Figure 3 b; Sup. Figure 12). Figure 3 b shows an example slice in which CB (green) correlates with Purkinje cells and their dendrites, GFAP (red) with astrocytes around blood vessels, VGluT1 (cyan) with parallel fiber boutons in the molecular layer as well as mossy fiber terminals in the granule layer, Kv1.2 (yellow) with pinceau surrounding Purkinje cell axons, and PV (magenta) with both Purkinje cells and molecular interneurons.

In order to facilitate visualization of the vCLEM dataset, we imported it into Neuroglancer ^52^ where each fluorescence channel and the EM channel can be visualized separately or simultaneously, and navigation through slices and different resolution levels can be carried out easily.

### 3D reconstruction of cells and subcellular structures identified by scFv labeling in the cerebellar cortex

An important question is whether the EM data, with six-color immunolabeling superimposed, is of sufficient quality to be successfully segmented by automatic means. Two different methods of automatic segmentation were successful. We used a flood-filling network ^53^ pretrained on a different dataset ^54^ at either 32 nm or 16 nm and without any new ground truth (Sup. Figure13 a, b). We also used another method developed in our lab ^55, 56^ to generate high-quality 2D membrane predictions, as well as 2D segmentation at 8 nm resolution (Sup. Figure13 c, d) following several rounds of training with manually generated ground truth. The success of the automatic segmentation indicates that the ultrastructure is not compromised by the detergent-free scFv immunofluorescence labeling protocol.

We then analyzed the CLEM data by reconstructing the EM ultrastructure at sites labeled with each of the five molecular markers (CB, GFAP, PV, Kv1.2, and VGluT1).

Calbindin (CB): In the cerebellar cortex, CB is expressed by Purkinje cells and some Golgi cells in the granule layer ^42, 57, 58^. In the vCLEM dataset, labeling of the anti-CB scFv (green fluorescence signal in Figure 4 a; Sup. Figure 14) corresponded to Purkinje cell bodies (Figure 4 a, b), a few likely large Golgi cells in the upper granule cell layer ^59^ (Sup. Figure 14 b, c), and heavily myelinated axons that presumably arose from Purkinje cells (Sup. Figure 14 a, inset 2). 3D reconstructions of one of the labeled Purkinje cells (Figure 4 c; Sup. Figure 14 a) and one of the labeled Golgi cells (Sup. Figure 14 d) showed that their morphologies were similar to previous Golgi/ultrastructural studies of rat cerebellum ^59^. The synaptic inputs from parallel fibers onto Purkinje cells are also obvious (Figure 4 c, d). In addition, we found that the dendritic trees of Purkinje cells in this region (Crus I) were not perpendicular to the rostral-caudal axis but intersected at the axis at an angle of around 55° (Sup. Figure 14 a, inset 1).

**Figure 4.**
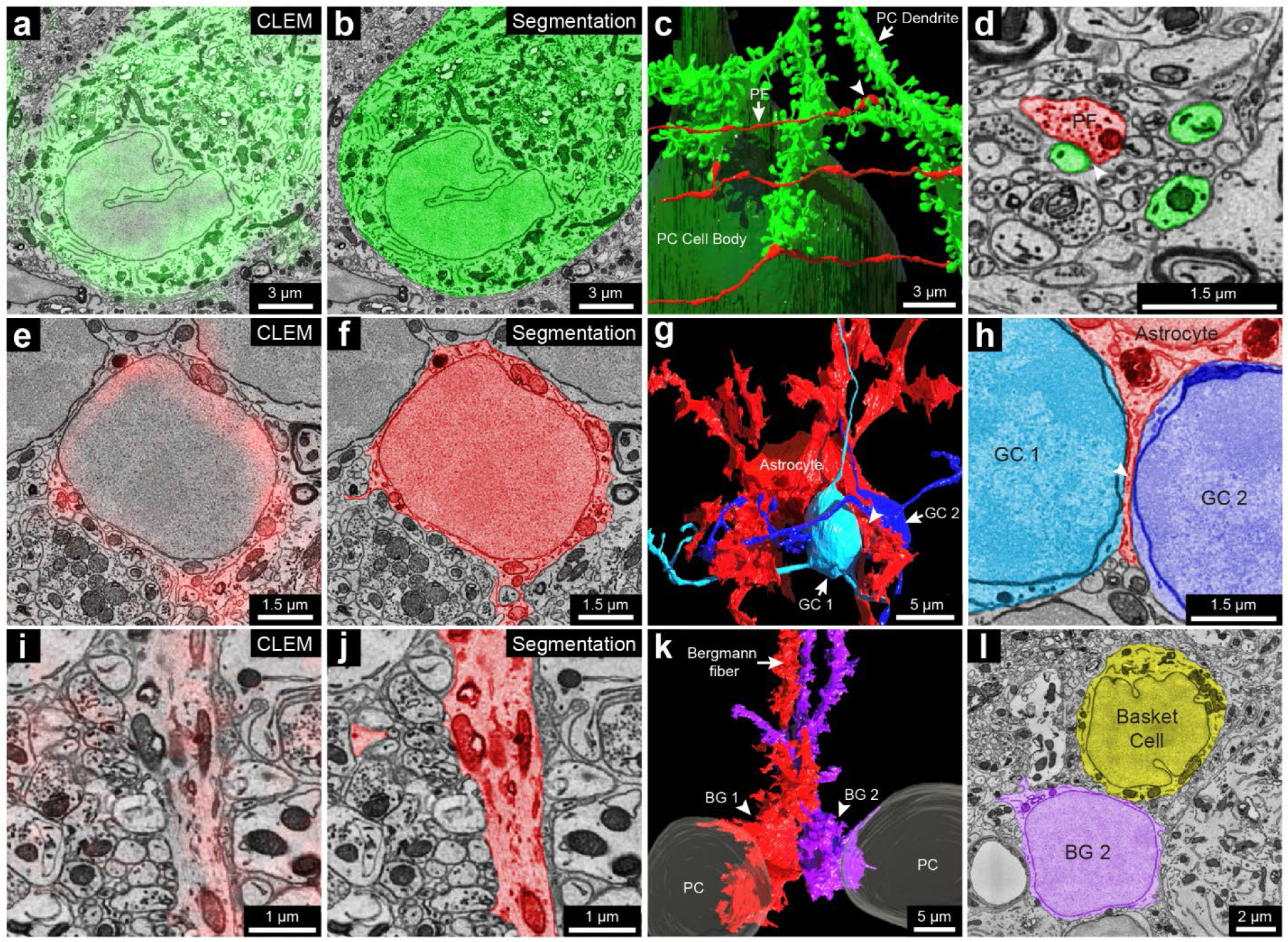
3D reconstruction of cells labeled by the calbindin-specific and the GFAP-specific scFv probes. **a**, 2D CLEM image showing the fluorescence signal (green) of the calbindin-specific scFv probe overlaps with the cell body of a Purkinje cell. **b**, EM image showing the 2D segmentation (green) of the calbindin positive Purkinje cell based on labeling shown in **a**. **c**, 3D reconstruction of the Purkinje cell labeled in **a**. The cell body is labeled in dark green. A dendritic branch is labeled in light green. Three parallel fibers are shown that made three synapses onto three spine heads of the dendritic branch. Arrowhead, a synapse made by a parallel fiber onto a spine head. PC, Purkinje cell; PF, parallel fiber. **d**, EM image showing the 2D segmentation of the synapse labeled by the arrowhead in **c** between a parallel fiber (red) and a spine head of the dendritic branch (green). **e**, 2D CLEM image showing the fluorescence signals (red) of the GFAP-specific scFv probe overlaps with the cell body of a velate astrocyte in the granule cell layer. **f**, EM image showing the 2D segmentation (red) of the cell body of the velate astrocyte in **e**. **g**, 3D reconstruction of the velate astrocyte (red) labeled in **e** and two nearby granule cells (light and dark blue). The velate astrocyte extended a veil-like glial process (arrowhead) between the two granule cells. GC, granule cell. **h**, EM image showing the 2D segmentation of the veil-like glial process (arrowhead) between the cell bodies of two granule cells. **i**, 2D CLEM image showing the fluorescence signals (red) of the GFAP-specific scFv probe overlaps with a Bergmann fiber. **j**, EM image showing the 2D segmentation (red) of the Bergmann fiber labeled in **i**. **k**, 3D reconstruction of two Bergmann glial cells (BG1 and BG2) traced back from their Bergmann fibers labeled by the GFAP-specific scFv probe (e.g. in **i**). BG, Bergmann glial cell. **l**, EM image showing the 2D segmentation of the cell body of BG2 and a nearby basket cell. Note the lack of infoldings in BG2’s nuclear membrane compared to the basket cell.

GFAP: This protein is expressed by astrocytes and Bergmann glial cells whose cell bodies reside in the Purkinje layer and whose processes (also known as Bergmann fibers) extend into the molecular layer ^60, 61^. As expected, labeling seen with the anti-GFAP scFv (red fluorescence signal in Figure 4 e, i) corresponded to the cell bodies and processes of granule layer astrocytes (Figure 4 e, f) and the vertical Bergmann fibers in the molecular layer (Figure 4 i, j). Although the cell bodies of granule layer astrocytes have a similar appearance to those of granule cells in EM micrographs (Figure 4 f; Sup. Figure 15 a, insets 1,2), 3D reconstruction of one of the labeled cells shows the typical morphology of a velate astrocyte ^59^ that is different from the structure of nearby granule cells, despite their similar cell body ultrastructure (Figure 4 g; Sup. Figure 15 a). The reconstructed velate astrocyte extended a thin, veil-like glial process between two granule cells (Figure 4 g, h). GFAP labeling was also seen in the processes of Bergmann glia but not in their cell bodies (Figure 4 i; Sup. Figure 15 b, insets). Two Bergmann glia cell bodies were found by tracing back from their labeled processes (Figure 4 i; Sup. Figure 15 b). These cell bodies were adjacent and between two Purkinje cells (Figure 4 k; Sup. Figure 15 b). Although they were similar in size to nearby basket cells, they were distinguishable by virtue of the lack of infoldings in their nuclear membranes (Figure 4 l).

Parvalbumin (PV): This protein is expressed by molecular layer interneurons (MLIs), Purkinje cells and Purkinje layer interneurons ^42, 62^. The labeling we observed (magenta fluorescence signal in Figure 5 a; Sup. Figure 16 a, b) in the molecular layer co-registered with MLIs (Figure 4 b; Sup. Figure 16 c, d) that have in-folded nuclear membranes. Among PV positive MLIs (n = 22) that we partially reconstructed we chose three for full reconstruction. These neurons were located at three different levels of the molecular layer (Figure 4 c) and exhibited different morphologies as has been previously described ^62^.

**Figure 5.**
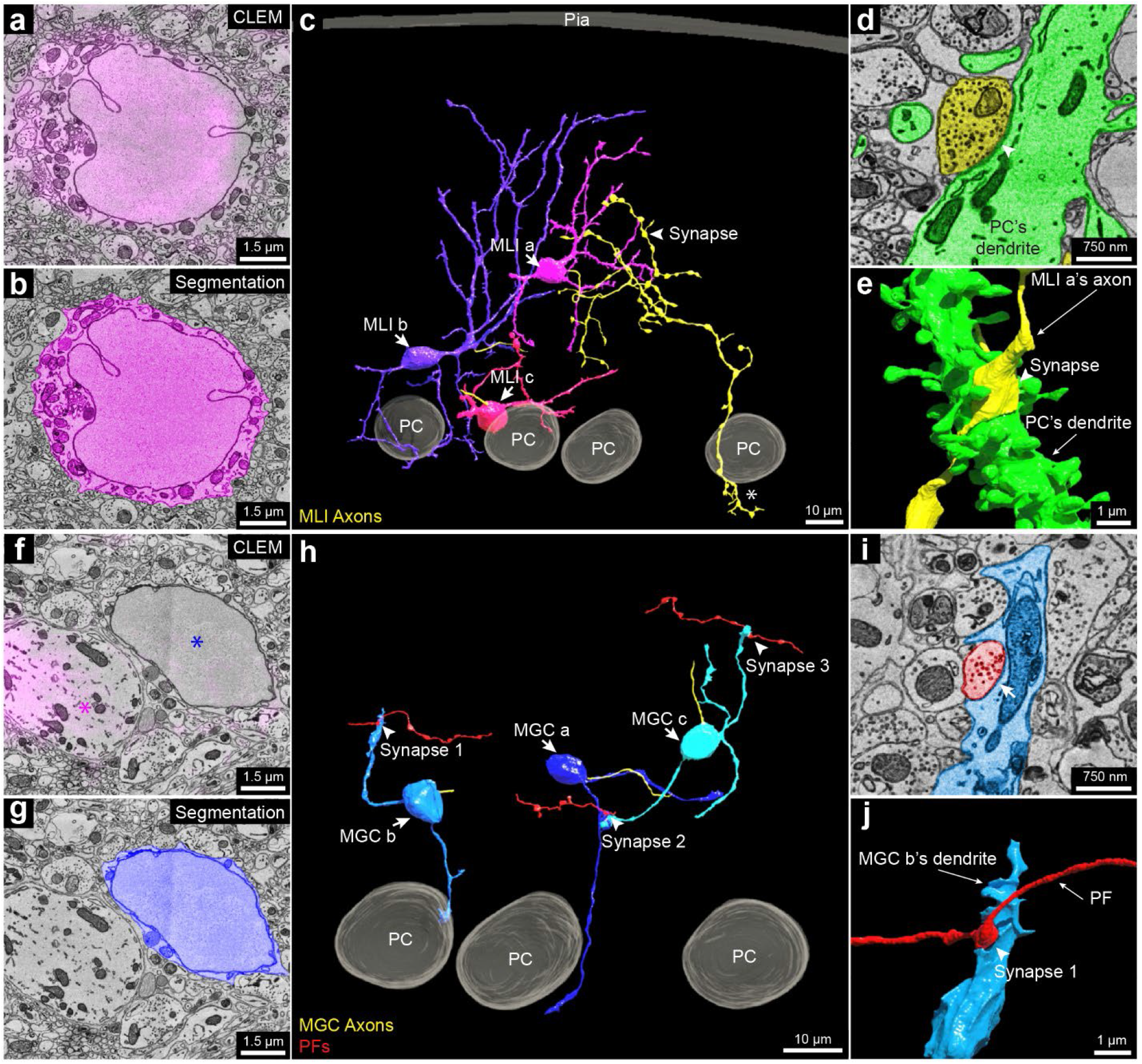
3D reconstruction of the molecular layer interneurons and granule cells distinguished by PV-specific scFv probes. **a**, 2D CLEM image showing the fluorescence signals (magenta) of the parvalbumin-specific scFv probe overlap a cell body of a molecular layer interneuron (MLI, also known as MLI a in **c**). **b**, EM image showing the 2D segmentation (magenta) of the MLI in **a**. **c**, 3D reconstruction MLI a, MLI b, and MLI c, showing their positions relative to the pia and the Purkinje cell layer. Their axons are labeled in yellow. MLI a, the interneuron that is farthest from the Purkinje cell layer axon branched extensively in the volume and innervated a Purkinje cell’s dendritic shaft at the arrowhead. This same axon was also part of the pinceau structure that surrounds a different Purkinje cell’s axon initial segment (asterisk, also see Sup. Figure 16 e). Based on these features, this interneuron appears to be a deep axon stellate cell ^59^. **d**, EM image showing the 2D segmentation of the synapse labeled by the arrowhead in **c** between MLI a’s axon (yellow) and a dendrite of a Purkinje cell (green). Arrowhead points to the location of the synapse. **e**, 3D reconstruction of the synapse labeled by the arrowhead in **d**. **f**, 2D CLEM image showing the fluorescence signal (magenta) of the parvalbumin-specific scFv which overlaps dendrites of a Purkinje cell (magenta asterisk), but does not label a molecular layer granule cell (blue asterisk). **g**, Same EM image as **f** showing the 2D segmentation (dark blue) of the cell body of a molecular layer granule cell (MGC). **h,** 3D reconstructions of MGC a, MGC b, and MGC c, with relationship to three Purkinje cells (also shown in the right part of **c**). Unlike granule cells with cell bodies in the granule cell layer, these MGCs received synapses from parallel fibers. Three such synapses between parallel fibers (red) and the three MGCs are labeled. **i**, EM image showing the 2D segmentation of synapse 1 in **h**. **j**, 3D reconstruction of synapse 1 labeled by the arrowhead in **i**.

The PV-positive labeling allowed examination of a set of neurons in the molecular layer that are PV-negative (Figure 5 f, g; Sup. Figure 16 f, g). All of these neurons (n=7) showed morphological features consistent with them being a rarely studied type of neuron, known as molecular layer granule cells (MGCs) ^59, 63, 64^. MGCs are sometimes regarded as ectopic granule cells that did not migrate to the granule cell layer during development ^59^. A recent study ^65^ showed that they are in fact relatively common and that they behave electrophysiologically like typical granule cells located in the granule cell layer. The MCG/MLI ratio of our dataset was 32%, which is similar to a previous quantification for the posterior lobe of the cerebellum from which our sample was derived ^65^. As mentioned, these neurons lack PV labeling in their cell bodies (Figure 5 f; Sup. Figure 16 f, g). Their cell bodies share ultrastructural features with typical granule cells. For example, they had round nuclear membranes without infoldings and only a small rim of cytoplasm surrounding the nucleus (Figure 5 g; Sup. Figure 16 h, i). Reconstruction of three MGCs showed that their 3D morphology was also similar to that of granule cells (Figure 5 h). They possessed several short dendrites, each forming a loosely organized glomerulus (Sup. Figure 16 j, insets 1, 2). At least sometimes the glomeruli were associated with mossy fiber collaterals that reached the Purkinje cell layer or slightly above (Sup. Figure 16 j, insets 3, 4). This is expected based on previous descriptions of mossy fibers ^59^. Surprisingly, the MGC dendrites were also innervated by parallel fibers (Figure 5 h to j). This has not been reported previously and is never the case for typical granule cells.

Kv1.2: This protein is a member of the voltage-gated potassium channel subfamily A ^66^, and shows high expression in the axonal terminals of the MLIs that form pinceau structures around Purkinje cell axon initial segments (AIS), as indicated by the strong yellow fluorescence signal in Figure 6 a and the reconstruction shown in Figure 6 b, c ^67, 68^. These axons are of different caliber (Sup. Figure 17 a-1 to a-7), and none are downstream branches of the same axon, suggesting that they each come from a different MLI cell. Kv1.2 also has been reported to be present in small puncta in the granule layer at juxtaparanodes of axons of unknown identity ^69^. We examined 22 sites of juxtaparanodal puncta in 22 axons in the granule cell layer. Consistent with previous reports ^66, 69^, we found 15 pairs of puncta that corresponded to the juxtaparanodal portion of nodes of Ranvier (Figure 6 d to j). In addition, we found 2 sites with three puncta that corresponded to the juxtaparanodal portion of nodes of Ranvier that are at branch points (Sup. Figure 17 b); Finally, we found 5 sites with a single punctum, which corresponded to the final myelinated site of an axon where myelination ended, a hemi-node of Ranvier (Sup. Figure 17 c). From the ssEM, we reconstructed these labeled axons. Six out of 22 had *en passant* or terminal axonal arborizations in the granule cell layer innervating granule cells (Figure 6 k; Sup. Figure 17c), indicating that they are mossy fibers. The remainder of the axons which had a similar appearance ultrastructurally but ran out of our EM volume prevented an unambiguous identification of their type, but they could also be mossy fibers. None of these axons were calbindin-positive. Axons that we suspect are primary Purkinje cell axons or their branches (because they were calbindin positive; Sup. Figure 14 a, inset 2) did not show Kv1.2 labeling at their juxtaparanodes. We thus conclude that at least a portion of the axons with Kv1.2 positive juxtaparanodes are mossy fibers, but not Purkinje cell axons.

**Figure 6.**
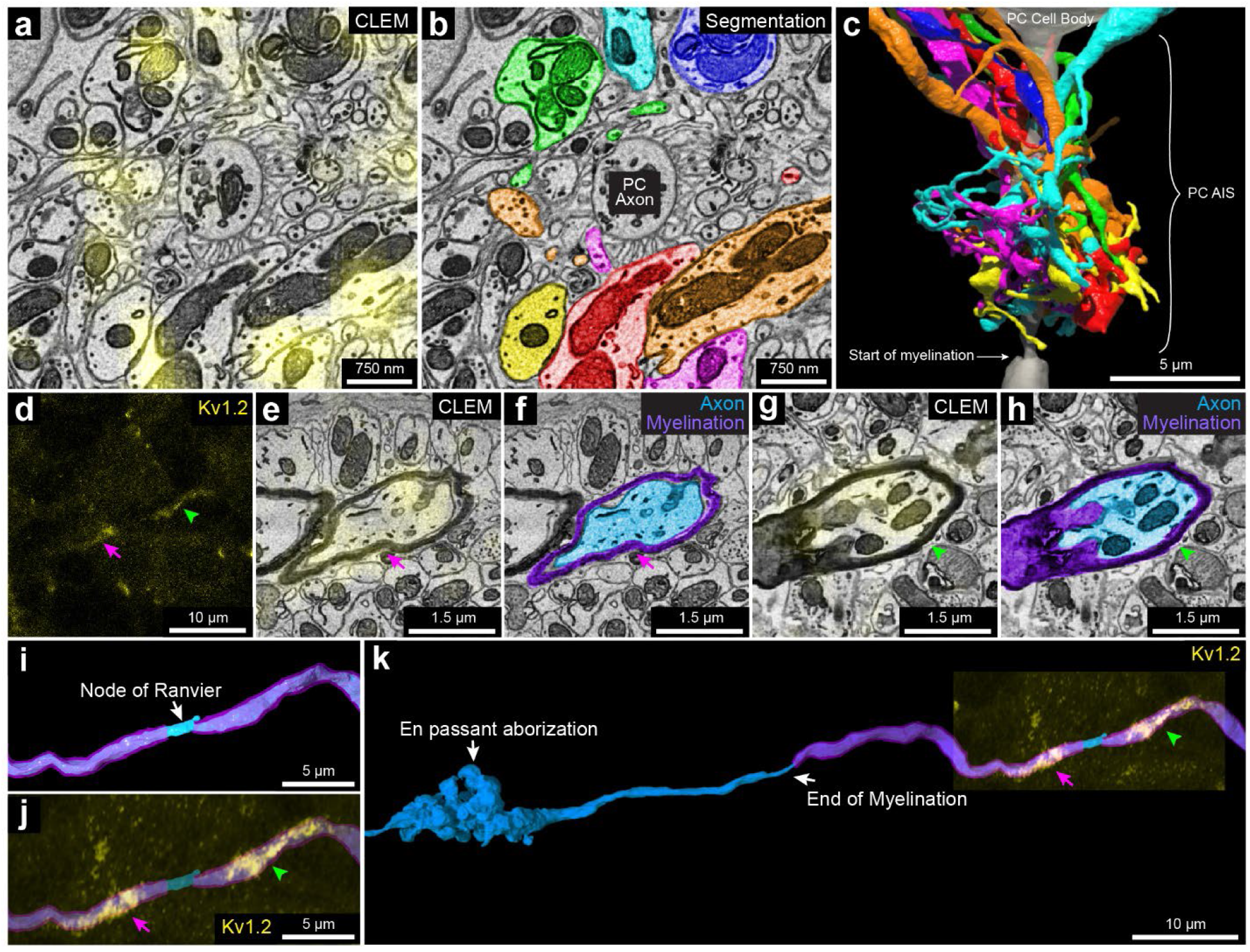
3D reconstruction of subcellular components with the Kv1.2-specific scFv probe in cerebellum. **a**, 2D CLEM image showing the fluorescence signals (yellow) of the Kv1.2-specific scFv probe overlaps with axonal terminals of MLIs at a pinceau structure surrounding the axon initial segment of a Purkinje cell. **b**, EM image showing the 2D segmentations of some (n=7) of the axon terminals labeled in **a**. **c**, 3D reconstruction of the seven axonal terminals in **b** (see Sup. Figure 17 a for the morphology of these axonal terminals). **d**, A pair of juxtaparanodal punctate labeling sites of the Kv1.2-specific scFv probe in the granule cell layer within the confocal image stack. The magenta arrow and green arrowhead show each side of the juxtaparanodal labeling. **e**, 2D CLEM image showing one side of the juxtaparanodal punctate labeling (yellow, magenta arrow) of the axon in **d**. **f**, Shows the 2D segmentation of the axon in **e**. **g**, 2D CLEM image showing another side of the juxtaparanodal punctate labeling (yellow, green arrowhead) of the Kv 1.2-specific scFv probe at the other side of the juxtaparanodal segment of the same node of Ranvier. **h**, EM image showing the 2D segmentation of the axon in **g**. **i**, 3D reconstruction of the node of Ranvier of the axon labeled in **d**-**h** (myelination labeled in purple). **j,** The juxtaparanodal labeling by the Kv1.2-specific probe in **d** overlaid onto the reconstructed node of Ranvier. **k**, More extended 3D reconstruction of the axon in **i** shows that it had an en passant arborization and another terminal arborization (not shown) in the granule cell layer, that appeared to be mossy fiber synapses.

VGluT1: VGluT1 is a member of the vesicular glutamate transporter family that is expressed by axonal terminals ^70^. In the cerebellar cortex, VgluT1 is highly expressed in the boutons of parallel fibers and in a subgroup of mossy fiber arborizations ^70, 71^. In Crus 1, both in a previous study ^71^ and in our immunofluorescence images, there are at least three types of mossy fiber arborizations in the granule cell layer: VGluT1 positive, VGluT2 positive, VGluT1/2 double positive terminals (Figure 7 a, c; Sup. Figure 18 a, b-1 to b-3). In our vCLEM dataset, the labeling of anti-VGluT1 scFv in the granule cell layer (cyan fluorescence signal in Figure 7 a) does not label all mossy fibers (Figure 7 c). We classified the mossy fiber arborizations based on whether they were VGluT1 positive (Figure 7 a, b) or VGluT1 negative (Figure 7 c, d). While we can’t further subdivide the VGluT1 positive from the VGluT1/2 double positive types, we surmise that the VGluT1 negative terminals are VGluT 2 positive. To see if these two categories had ultrastructural differences, we randomly chose ten VGluT1 positive and ten VGluT1 negative mossy fiber terminals and reconstructed them. The 3D morphologies of both types were similar (Figure 7 e to g). However, the mean volumes of VGluT1 positive terminals were significantly larger than VGluT1 negative terminals (p = 0.0191; p < 0.05; t=2.575, df=18) (Figure 7 h) (see Sup. Table 4 and 5 for the volumes of each terminal). We also performed automatic detection of synaptic vesicles (Sup. Figure 18 c, e) and mitochondria in these terminals (Sup. Figure 18 d, f) using machine learning algorithms (see Sup. Table 4 and 5 for the vesicle number, vesicle density, mitochondrial volume, and the mitochondria per terminal volume of each terminal). The mean value of the number of synaptic vesicles per terminal was significantly higher in VGluT1 positive terminals than in VGluT1 negative terminals (p = 0.0050; p < 0.01; t=3.200, df=18) (Figure 7 i). The mean value of the synaptic density per terminal trended higher in VGluT1 positive terminals than in VGluT1 negative terminals (p = 0.0705; p < 0.1; t=1.922, df=18) (Figure 7 j). The mean value of the mitochondria per terminal volume was not significantly different (p = 0.1285; p > 0.1; t=1.593, df=18) (Figure 7 k). These results suggest that these two types of mossy fiber terminals have distinct EM ultrastructure. They may originate from different brain regions and have different electrophysiological properties. However, we found a number of granule cells that were innervated by both VGluT1 positive and VGluT1 negative mossy fibers.

**Figure 7.**
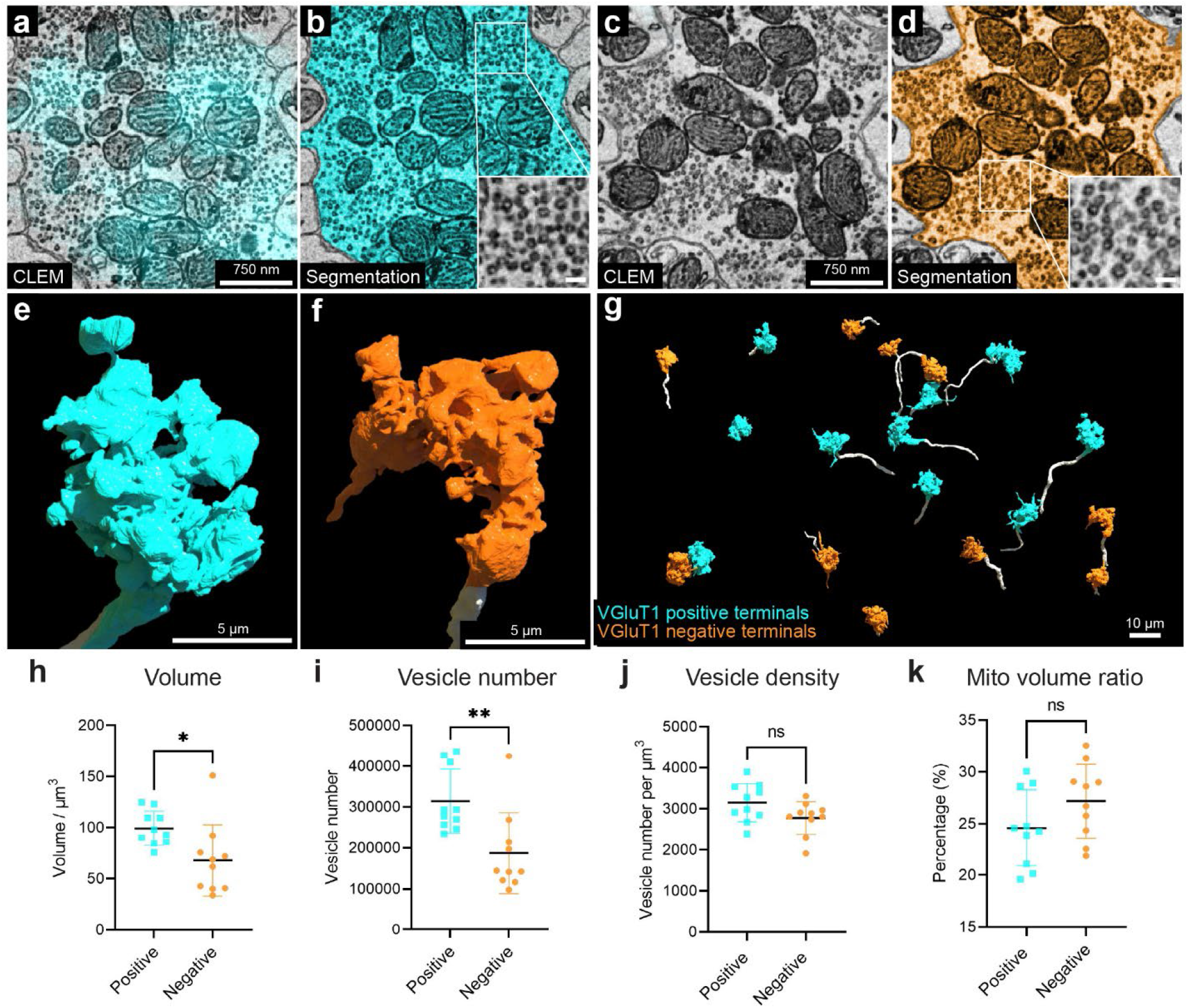
3D reconstruction and analysis of VGluT1 positive and negative mossy fiber terminals based on labeling with the VGluT1-specific scFv probe. **a**, 2D CLEM image showing the fluorescence signals (cyan) of the VGluT1-specific scFv probe overlaps with a mossy fiber terminal. **b**, EM image showing the 2D segmentation (cyan) of the mossy fiber terminal labeled in **a**. The enlarged inset shows the morphology of synaptic vesicles in this terminal. **c**, 2D CLEM image showing a mossy fiber terminal lacking VGluT1 fluorescence signal. **d**, EM image showing the 2D segmentation (orange) of the mossy fiber terminal labeled in **c**. The enlarged inset shows the morphology of synaptic vesicles in this terminal. **e**, 3D reconstruction of the VGLUT1 positive mossy fiber terminal (cyan) in **b**. The axon leading to the terminal arises from the bottom of the panel. **f**, 3D reconstruction of the VGluT1 negative mossy fiber terminal (orange) in **c**. The axon leading to the terminal arises from the bottom of the panel. **g**, 3D reconstruction of ten VGluT1 positive mossy fiber terminals (cyan) and ten VGluT1 negative mossy fiber terminals (orange). The axons are labeled in white. **h**, volume, **i**, vesicle number, **j**, versicle density, **k**, mitochondria volume ratio per terminal volume were measured for each of the 10 VGluT1 positive terminals, cyan, and the 10 VGluT1 negative terminals, orange. Mean values with SD were shown on each graph. Mito, mitochondria.

## Discussion

Here we report a technique that visualizes multiple molecular labels superimposed on an electron microscopic volume, using detergent-free scFv-based immunofluorescence. We used this method to study a volume of cerebellum labeled with five scFv probes, acquired by serial section electron microscopy. This vCLEM dataset provided several new insights relevant for the cytology of cerebellar cortex. We deposited the expression plasmids of eight scFv probes in an open access repository ^72^ (see Table 1 for the links at Addgene). All of these probes possess a sortase tag to allow straightforward attachment of fluorescent or other labels for use in vCLEM studies or other applications.

Expanding this technique to more probes will require acquiring a larger collection of functional scFv probes to include additional commonly used molecular markers in bioscience. The scFv probes we generated were all based on mAbs from NeuroMab ^36^. This facility has developed thousands of mAbs for neuroscience research, a subset of which have been converted into recombinant form ^29^. Consistent with the overall track record of converting mAbs into scFvs ^73^, current efforts to systematically convert NeuroMab mAbs into scFvs that retain the binding characteristics of the progenitor mAb is ∼ 50%. All of the scFvs described here label brain tissue. To generate more scFvs, several strategies present themselves. First, *de novo* amino acid sequencing of a mAb can establish the sequences of the V_H_ and V_L_ domains of other commonly used mAbs or even individual components of polyclonal Abs for subsequent conversion into scFvs ^74, 75^. Second, artificial protein evolution using phage-display methods has been used to modify the structure of failed scFvs to render them functional ^76^. Third, rational protein design via point-mutation or CDR-grafting has been used to improve the performance of failed scFvs ^77, 78^. Finally, purely *in vitro* approaches using phage display or other methods can be used to screen naïve or immune scFv libraries to develop novel scFvs ^79^. We therefore anticipate that many more functional scFv probes will become available, in particular through open access sources.

Multiplexability, i.e., multiple labels with different colors, can make vCLEM more cost-effective and more powerful. More resolvable color labels in a specimen means that more molecular information can be extracted without the need of multiple samples. Moreover, combinatorial labels may parse more cell types than possible with only a single label at a time. For example, in the cerebellum, cells that are CB- and PV-positive are Purkinje cells, whereas PV-positive, but CB-negative cells, are molecular layer interneurons. This same combinatorial approach will likely be useful in parsing various subtypes of interneurons in the cerebral cortex.

With spectral unmixing, we were able to acquire six-color fluorescent image volumes from samples immunolabeled with scFv probes. However, six-color images are not the upper limit of how many colors can be disambiguated. With the growing list of functional scFvs, optimized algorithms of linear unmixing can achieve super-multiplexed fluorescence imaging with up to 45 colors ^15^. It is possible that scFvs could also be conjugated with narrow-peaked Raman imaging dyes to achieve multiplexed Raman vibrational imaging with 24 or more colors ^80^. In addition, as our technique is based on immunolabeling, it can be easily combined with other types of functional vCLEM. For example, scFv-enabled immunofluorescence can be applied to samples that have been functionally imaged with calcium or voltage indicators, so that the molecular identities of the neurons whose functional properties have been analyzed can also be determined. This advantage can greatly expand the knowledge of neural networks examined by functional live imaging.

We used confocal microscopy to acquire fluorescence volumes of scFv-labeled samples because of the power of spectral unmixing to separate multicolor fluorescence. The superior penetration ability of scFvs as demonstrated in this study already exceeds the imaging depth of confocal microscopy of uncleared tissue. Our approach generates usable vCLEM volumes up to a depth of 120 μm. Even deeper imaging may be possible with spectral unmixing techniques for two-photon and light sheet microscopies ^81, 82^.

With the rapid progress in ssEM, very large datasets (∼1 mm^3^) ^54, 83^ are being generated. However, the generation of vCLEM datasets of such sizes remains technically problematic. However, the use of scFv labels, once associated with electron microscopy ultrastructure, may circumvent this problem. For example, the cerebellar cortex dataset showed that despite their similar location, the cell bodies of Bergmann glia, MLIs and MGCs were distinct (Figure 4 l; Figure 5 a, b, f, g) and could conceivably be used to train a classifier for determination of cell type. Indeed, using machine learning we could differentiate two molecularly defined mossy terminal types, despite them having the same neurotransmitter and being functionally and gross-anatomically similar (Figure 7). More recent machine learning algorithms like “SegCLR” ^84^ use data from small volumes and then identified cell types in large volumes. We believe that the growing number of probes for vCLEM, combined with machine learning, will allow the classification of cellular and subcellular molecular types in unlabeled electron microscopy images.

## Methods

### Animals

Animals used in the study were adult C58BL/6J mice (Jackson Laboratory). All experiments using animals were conducted according to US National Institutes of Health guidelines and approved by the Committee on Animal Care at Harvard University.

### ScFv production

The amino acid sequence of scFv was designed by linking the variable domains of the heavy chain and the light chain of a mAb with a (GGGGS GGGGS GGGGS) linker (3x linker), a (GGGGS GGGGS GGGGS GGGS) linker (4x linker 1), or a (GGGGS GGGGS GGGGS GGGGS) linker (4x linker 2). A signal peptide (MDWTWRILFLVAAATGAHS or MGWSCIILFLVATATGVHS) was added to the N’-terminus. A sortase tag (SLPETGG) and a 6 x His tag was added to the C’-terminus. The amino acid sequence was reverse translated into a DNA sequence with codon optimization for human cells. The DNA sequence was synthesized and cloned into pcDNA 3.1 or pcDNA 3.4 vectors.

Expi 293 cells (ThermoFisher) were used for scFv expression. The cells were maintained in Expi 293 Expression Medium (ThermoFisher) and passaged every other day. After two rounds of passage, transfection of the scFv expression plasmids was carried out using 293fectin (ThermoFisher) following vendor’s protocol. For 100 ml culture with 1x 10^8^ cells, we used 100 µg plasmid DNA and 200 µl 293fectin. Cells were incubated for another seven days, after which the culture supernatant was collected. The supernatant was incubated with Ni-NTA agarose (ThermoFisher). After washing with Tris buffer (50 mM Tris, 150 mM NaCl, pH = 7.5∼8.0), the bound proteins were eluted with the elution buffer (250 mM imidazole, 50 mM Tris, 150 mM NaCl, pH = 7.5∼8.0). The eluted proteins were loaded into Amicon® Ultra-15 centrifugal filter unit (Millipore Sigma), washed with Tris buffer, and then buffer exchanged into Tris buffer with 15% glycerol by centrifugation at 4000 rpm at 4 °C. The concentration of the purified protein was determined by A280. The purified protein was stored at -20 °C.

### Sortase reaction

The GGGC peptide was synthesized at BIOMATIK. The GGGC peptides were dissolved in 1 x PBS (Sigma-Aldrich) at the concentration of 20 mg/ml. The fluorescent dyes in maleimide form (ThermoFisher) were dissolved in DMSO at the concentration of 20 mM. The dissolved peptide and fluorescent dyes were mixed at the molar ratio of 12:1 and then incubated with shaking at 800 rpm at 4 °C overnight. The GGGC-dye conjugates were purified by HPLC at the Koch Institute at MIT. Identity of the purified conjugates were confirmed by mass spectroscopy, lyophilized, and then re-dissolved in ddH_2_O at the concentration of 4 mM.

The sortase reaction was performed as described in ^85^. In brief, GGGC-dye conjugates (500 uM) and sortase tag-containing scFvs (100 uM) were mixed in the sortase reaction buffer (50 mM Tris-HCl, 150 mM NaCl, 10 mM CaCl_2_, pH = 7.5∼8.0), followed by adding sortase A pentamutant that has a 6 x His tag (2.5 uM). The mixture was incubated with shaking at 500 rpm at 12 °C for 3 h. Then Ni-NTA agarose was added into the mixture, followed by an incubation shaking at 500 rpm at RT for 20 min to remove sortase and any leftover GGGC-dye conjugates. After the incubation, the mixture was loaded into a microcentrifuge spin column (USA Scientific) and centrifuged briefly to remove the Ni-NTA agarose. The filtered solution that contains the scFv-dye conjugates was loaded into Amicon® Ultra-4 centrifugal filter unit (Millipore Sigma), washed with Tris buffer and then buffer exchanged into Tris buffer with 15% glycerol by centrifugation at 4000 rpm at 4 °C. The concentration of the purified scFv-dye conjugates was determined by A280. The purified scFv-dye conjugates were stored at -20 °C.

### Perfusion and fixation

The mouse was anesthetized by isoflurane until there was no toe-pinch reflex. Mice were then transcardially perfused with aCSF (125 mM NaCl, 26 mM NaHCO_3_, 1.25 mM NaH_2_PO_4_, 2.5 mM KCl, 26 mM glucose, 1 mM MgCl_2_ and 2 mM CaCl_2_ (all chemicals were from Sigma-Aldrich) at the flow rate of 10 ml/min for 2 min to remove blood, followed with 4% paraformaldehyde (Electron Microscopy Sciences), 0.1% glutaraldehyde (Electron Microscopy Sciences) in 1 x PBS for 3 min for fixation. Brains were dissected and then post-fixed in the same fixative on a rotator overnight at 4°C. Brains were sectioned into 50-μm or 120-μm coronal sections using a Leica VT1000 S vibratome and stored in the same fixative at 4°C.

For ECS-preserving perfusion, the detailed protocol was described in ^27^. In brief, mice were anesthetized by isoflurane, transcardially perfused with aCSF at the flow rate of 10 ml/min for 2 min to remove blood, followed with 15 w/v% mannitol (Sigma-Aldrich) aCSF solution for 1 min, 4 w/v% mannitol aCSF solution for 5 min, and 4 w/v% mannitol, 4% paraformaldehyde in 1 x PBS for 5 min. Brains were dissected out and then post-fixed in the same fixative for 3 h on a rotator at 4 °C. Brains were sectioned into 50-µm coronal sections using a Leica VT1000 S vibratome, and then store in in 1 x PBS at 4 °C.

### Immunofluorescence

Detergent-free immunofluorescence labeling was performed with scFv probes or nanobody probes (see Sup. Table 1 for the scFv and nanobody probes used in this study). The detailed protocol was described in ^23^. In brief, 50-µm or 120-µm coronal sections were transferred into 3.7 ml shell vials (Electron Microscopy Sciences) with 1 ml 1 x PBS. The sections were first washed with 1 x PBS for 3 x 10 min, and then blocked in glycine blocking buffer (0.1 M glycine, 0.05% NaN_3_ in 1 x PBS) for 1 h on a rotator at 4 °C. The labeling solution was prepared by diluting scFv-dye conjugates or nanobody-dye conjugates in glycine blocking buffer (see Sup. Table 1 for the final concentration of each scFv and nanobody probe). The labeling solution and any remaining steps were protected from light. The sections were then incubated with the labeling solution on a rotator at 4 °C. 50-µm sections were incubated for 3 days. 120-µm sections were incubated for 7 days. After the incubation, sections were washed with 1 x PBS for 3 x 10 min, and then stained with Hoechst 33342 (Invitrogen, diluted 1:5000 in 1 x PBS) for 1 h on a rotator at 4 °C. Sections were washed with 1 x PBS for 3 x 10 min, and then mounted onto glass slides (VWR).

Immunofluorescence labeling with detergent was performed with primary antibodies plus secondary antibodies for single labeling (for the validation of scFv probes versus mAbs; see Sup. Figure 2-4), or with primary antibodies plus secondary antibodies and scFv probes for double labeling (for the double labeling experiment of VGluT1 and VGuT2; see Sup. Figure 18) (see Sup. Table 2 for the primary antibodies and Sup. Table 3 for the secondary antibodies used in this study). The detailed protocol is available on the NeuroMab website ^41^. In brief, 50-µm coronal sections were transferred into 3.7 ml shell vials with 1 ml 1 x PBS. The sections were first washed with 1 x PBS for 3 x 10 min, and then blocked in vehicle (10% normal goat serum, 0.3% Triton X-100 in 1 x PBS) overnight on a rotator at 4 °C. Subsequent steps were protected from light. The sections were then incubated with the primary antibody solution or primary antibody solution plus scFv probes (for the experiment of the double labeling of VGluT1 and VGuT2; see Sup. Figure 18) (see Sup. Table 1, 2 for the dilution ratios of primary antibodies and the anti-VGluT 1 scFv probe used in this study;) for 2 days on a rotator at 4 °C. After the incubation, sections were washed with vehicle for 3 x 10 min, and then incubated with the secondary antibody solution (see Sup. Table 3 for the dilution ratios of secondary antibodies used in this study) overnight on a rotator at 4 °C. After the incubation, sections were washed with 1 x PBS for 3 x 10 min, and then stained with Hoechst (diluted 1:5000 in 1 x PBS) for 1 h on a rotator at 4 °C. Sections were washed with 1 x PBS for 3 x 10 min, and then mounted onto glass slides.

### Fluorescence confocal microscopy

50-µm sections were mounted in Vectashield H-1000 (Vector Laboratories) with a #1 coverslip (Electron Microscopy Sciences) on top sealed with clear nail polish. 120-µm sections (for CLEM) were mounted in 1 x PBS inside 120-um spacer (Invitrogen) with a #1 coverslip on top sealed with the spacer. 50-µm sections were imaged with a Zeiss LSM 880 confocal laser scanning microscope equipped with either a 20x/0.8 NA air-objective, or a 40x/1.1 NA water immersion objective, or a 63x/1.4 NA oil-immersion objective depending on the required optical resolution. Acquisition of double or triple color fluorescent images was done with appropriate band pass filters for the specific fluorescent dyes to avoid crosstalk.

For spectral imaging, the detailed protocol was described in ^48^.In brief, 120-µm sections immunolabeled by five scFv with different fluorescent dyes (Alexa fluor 488, 532, 594, 647, and 5-TAMRA) were imaged with a Zeiss LSM 880 confocal laser scanning microscope equipped with a 40x/1.1 NA water immersion objective. All fluorophores except Hoechst were excited simultaneously with 488, 560, 633 nm lasers and a 488/561/633 nm beam splitter. A z-stack of lambda stack images with the interval of 1 µm in z axis was acquired by an array of 32 Gallium Arsenide Phosphide (GaAsP) detectors in lambda mode at 9nm bins from 414 nm to 691 nm. Lambda stack images were acquired from each of the sections labeled by only one of the five scFvs with the same settings in lambda mode. Reference spectra for each fluorescent dye were extracted from each of these lambda stacks. Linear unmixing was performed on the z-stack of lambda stack images acquired from the multi-labelled section using the reference spectra in Zen Blue software (Zeiss). The Hoechst channel (labeling cell nuclei) of the multi-labelled section was acquired separately with a single 405 nm laser and appropriate band pass filters.

To test if the Hoechst channel could be combined into linear unmixing, we tried acquiring another z-stack of lambda stack images of another multi-labeled 120-µm section with simultaneous excitation with 405, 488, 560, 633 nm lasers, a 488/561/633 nm beam splitter and a 405 nm beam splitter. A lambda stack image was acquired from a section labeled by only Hoechst with the same settings in lambda mode. Linear unmixing was performed on the z-stack of lambda stack images of the multi-labelled section using the reference spectra (five fluorescent dyes plus Hoechst) in Zen Blue software (Zeiss). Several reference spectra for Hoechst extracted from different locations of pixels in Hoechst’s lambda stack image caused the Hoechst signals to be unmixed into the channel of Alexa 488 in error, but we did find one spectrum that generated good unmixing results (Sup. Figure 5).

The brightness, contrast, and gamma of all fluorescent images were adjusted. Fluorescence image volumes were projected to a single plane by maximum intensity for visualization in 2D.

### ScFv, nanobody, and antibody penetration experiment

Immunofluorescence with or without detergent by scFv probes, nanobody probes, primary antibodies with dyes directly conjugated, or with primary antibodies plus secondary antibodies was performed on 300-µm coronal sections from animals perfused with normal or ECS-preserving protocol. See above for detailed protocol of immunofluorescence. For the sections labeled with scFv probes, nanobody probes, primary antibodies with dyes directly conjugated, the incubation lasted for 7 days. For the sections labeled with the primary mAb for calbindin, the sections were incubated with the primary antibody solution (see Sup. Table 2 for the dilution ratio of this mAb) for 4 days on a rotator at 4 °C. After the incubation, sections were washed with vehicle for 3 x 10 min, and then incubated with the secondary antibody solution (see Sup. Table 3 for the dilution ratio of the secondary antibody) for 3 days on a rotator at 4 °C. After the incubation, sections were washed with 1 x PBS for 3 x 10 min. Then the sections were sliced into 50-µm sections lengthwise using a VT1000 S vibratome. The section from the middle was mounted onto glass slides, and then imaged by confocal microscopy to evaluate the penetration depth.

### EM preparation

After 120-µm sections were imaged, a small amount of 1 x PBS was added between the glass and coverslip to unseal the coverslip. Sections were transferred into 3.7 ml shell vials with 1 ml secondary fixative (2% PFA, 2,5% glutaraldehyde in 0.15M sodium cacodylate buffer with 4mM Ca^2+^ and 0.4 mM Mg^2+^) and incubated for at least one week on a rotator at 4 °C. A modified ROTO (Reduced Osmium-Thiocarbohydrazide-Osmium) protocol was used to stain the sections. All steps were carried out at room temperature (RT) except when specified. Sections were washed 3 x 10 min in 0.15 M sodium cacodylate buffer with 4 mM Ca^2+^ and 0.4 mM Mg^2+^, and then incubated in a solution containing 2 w/v% OsO_4_ (Electron Microscopy Sciences), 1.5% w/v potassium ferrocyanide (Sigma-Aldrich) and 0.15M cacodylate buffer inside 120-µm spacers (the top surfaces were not peeled) between a glass slide and a coverslip for 2 h with a change of the same solution after the first hour to prevents tissue deformation. Sections were incubated in the same solution in shell vials for another 1.5 h on a rotator. After washing 3 x 10 min in ddH_2_O, sections were incubated in a filtered aqueous solution containing 1 w/v% thiocarbohydrazide (Sigma-Aldrich) for 45 min. After washing 3 x 10 min in ddH_2_O, sections were incubated in an aqueous solution containing 2 w/v% OsO_4_ for 1.5 h. After washing 3 x 10 min in ddH_2_O, sections were incubated in a filtered aqueous solution containing 1 w/v% uranyl acetate (Electron Microscopy Sciences) overnight on a rotator protected from light. On the next day, after washing 3 x 10 min in ddH_2_O, sections were dehydrated in a series of graded (25, 50, 75, 100, 100, 100 v/v%, 10 min each) acetonitrile (Electron Microscopy Sciences) aqueous solutions. Afterward, sections were infiltrated with 25% LX-112 (Ladd Research) : acetonitrile for 1.5 h, 50% LX-112 : acetonitrile for 3 h, 75% LX-112 : acetonitrile overnight, 100% LX-112 for 12 h on a rotator. Then the sections were embedded in 100% LX-112 at the bottom of a flat bottom embedding capsule (Electron Microscopy Sciences) and cured in a 60 °C oven for two days.

### X-ray MicroCT scanning

X-ray micro-computed tomography (μCT) images of the resin-embedded sections were acquired using a Zeiss Xradia 510 Versa system and Zeiss’ Scout and Scan software. The imaging conditions include a projection pixel size of around 1.15 µm (pixel bin size of 1), 60 kV x-ray source, 30 s exposure per projection, and 3501 projections for 360 degrees of rotation. Each scan took approximately 33 hours. The Feldkamp, Davis, and Kress (FDK) algorithm reconstructed 3-dimensional volume maintains the projection pixel size of 1.15 µm resulting in a voxel size of 1.15 µm^3^. The Scout and Scan Control System Reconstructor software was used to convert the reconstructed files to .tiff files.

### EM imaging

The resin-embedded sections were cut into 30 nm serial ultrathin sections using automated tape-collecting ultramicrotome (ATUM) ^86^. Serial sections were collected onto carbon-coated and plasma-treated Kapton tape. The tape was cut into strips and affixed onto 150 mm silicon wafers (University Wafer).

A Zeiss Sigma scanning electron microscope was used to acquire overview images from the serial sections. Two overview images were taken per wafer, which were around 1.35 µm apart in z-axis. Typical imaging conditions are 8-kV landing energy, 1.2-nA beam current, 3-µs dwell time, 150-nm pixel size, and 4k x 4k images. The images were captured using a below-the-lens backscatter detector and Zeiss’ Atlas 5 software. The overview images were aligned using the “Linear stack alignment with SIFT” plugin in FIJI.

Prior to acquiring high-resolution images, the serial section sections on wafers were post-stained for 4 min with a 3% lead citrate solution. After staining, the sections were degassed for a minimum of 24 h at 1x10-6 Torr. A Zeiss MultiSEM 505 scanning electron microscope equipped with 61 electron beams was used to acquire high-resolution images from the serial sections. Images were collected using a 1.5-kV landing energy, 4-nm image pixel, and a 400-ns dwell time.

### High-resolution EM image processing

The preparation of the ssEM data before it could be segmented/analyzed includes two steps: affine stitching and elastic alignment. Affine stitching was performed on the raw data coming from the microscope. Firstly, SIFT features were extracted from the boundary area of each raw tile. Then all the neighborhood tiles in each section were matched together based on SIFT feature matching. Finally, a global optimization step made the matching results smooth. After each section was stitched well, an elastic alignment of all the sections was performed. A rough affine transformation between each mFov in section i to section i+1 and i+2 was estimated based on the SIFT features of blob-like objects detected in the sections. Then template matching, performed as a fine-grained matching for section i to section i+1 and i+2, was done on grid distributed image blocks in section i. At last, a global optimization on all the sections in the whole stack made the elastic transformation of mesh grid points on all the sections smooth.

The aligned stack was rendered at full resolution (4 x 4 x 30 nm) and each section was cut into 4k x 4k .png tiles, imported into VAST as a .vsvi file, and ingested into Neuroglancer for further analysis.

### Co-registration of fluorescence and EM volumes

The ssEM volume at a downsampled pixel size of 128 nm in the x and y plane and 480 nm in the z-axis was exported from the high-resolution .vsv file in VAST. The multi-color fluorescent image volume’s brightness and contrast were adjusted with the “Stack contrast adjustment” plugin in FIJI to compensate for signal decay in the z-axis. The pixel sizes in the x and y plane and in the z-axis were resampled to match those of the confocal fluorescence image volume. Both volumes were loaded into FIJI. With the BigWarp plugin, 188 landmark points were manually placed on corresponding sites of blood vessels, cell nuclei, cell bodies, and axons in the two volumes. The fluorescent image volume was 3D transformed by a thin plate spline interpolation based on the point correspondences. The transformed fluorescence volume was resampled in the z-axis and then imported into VAST as .vsv files to be overlaid with the ssEM volume at a downsampled pixel size of 128 nm in the x and y plane and 30 nm in the z-axis. The .vsv files were also ingested into Neuroglancer and resampled in the z-axis to be overlaid with the ssEM volume at a pixel size of 8 nm in the x and y plane and 30 nm in the z-axis.

### Automatic segmentation

The ssEM dataset was segmented in 3D using Flood-Filling Networks (FFNs) ^53^. The FFN segmentation model was trained at 32 x 32 x 30 or 16 x 16 x 30 nm resolution on the H01 dataset ^54^, and run here on CLAHE intensity normalized data ^87^ downsampled to match the trained model resolution. The resulting base supervoxels were assembled into larger per-cell segments via manual proofreading.

The ssEM dataset was also segmented in 2D at 8 x 8 x 30 nm resolution by a method developed in our lab. A description of this approach was given in ^55, 88^. In brief, a pre-trained algorithm was used to generate ground truth tiles of membrane predictions, which was corrected by a human annotator. Three rounds of ground truth correction were used to iteratively train deep neural network ^56^ with a UNET architecture ^89^. Neural network predictions were done on a commodity GPU using MATLAB connected to a VAST ^90^ instance, serving the EM images. In each round the model was applied on tiles randomly selected from the entire EM space and corrections were made to tiles that contained errors. Producing further training rounds halted when a sufficient accuracy was met. Finally, 2D segmentation using a region-growing algorithm was applied on the entire space according to the local minima of the membrane predictions.

### Mitochondria detection

We implemented an anisotropic U-Net architecture, incorporating a combination of 2D and 3D convolutions, to predict binary mitochondrial masks and instance maps similar to the U3D-BC approach in MitoEM ^91^. Given that the annotations for the new training dataset are confined to a select number of neurons rather than densely annotated, our model exclusively considers regions containing labeled neurons for loss computation. This strategy prevents the generation of empty predictions for valid mitochondria solely due to the absence of annotations within labeled neurons. To separate individual mitochondria, including those in close proximity, we employ marker-controlled watershed segmentation based on the network’s predictions. The segmentation system is implemented with the open-source PyTorch Connectomics deep learning toolbox ^92^.

### Vesicle segmentation and visualization

We first saturated segmented all vesicles on six images from a terminal, totaling around 8,400 instances. Then, we fine-tuned the Cytoplasm 2.0 model from Cellpose ^93, 94^ on these annotations. Specifically, as the model was pretrained on images with cells of a bigger size, we scaled up our image and segmentation maps of vesicles by 2x in both XY dimensions. For the 20 volumes of terminals, we obtained roughly 5 million vesicle instances.

### Statistical analysis

After the brightness of contrast of the layer of the fluorescent signals of VGluT1 were fixed at a value in Neuroglancer, a human annotator randomly picked 10 locations of VGluT1 positive mossy fiber terminals and 10 locations of VGluT1 negative mossy fiber terminals on slice 512 of the ssEM dataset. After 3D reconstruction of these terminals, the volume size of each terminal was generated using VAST. Mitochondria and synaptic detections (see above) were performed on segmentation of each terminal. None of these approaches were done blind to whether a terminal is VGluT1 positive or negative. Two-tailed, unpaired t-tests on volume, synaptic vesicle number, synaptic density, and mitochondria volume per terminal were performed in Prism-GraphPad.

## Acknowledgement

We thank the Harvard Center for Biological Imaging (RRID:SCR_018673) for infrastructure and support, D. Richardson at Harvard Center for Biological Imaging for advice on linear unmixing, the Biopolymers and Proteomics Core Facility at the Koch Institute at MIT for processing the peptide-fluorescent dye conjugates, the Bauer Core Facility at Harvard for infrastructure and support. This work was supported by NIH grants U19 NS104653, UG3 MH123386, P50 MH094271 (J. W. L.), U24 NS109113 (J. S. T.), and K99 MH128891 (X. L.); NSF grants NSF-CAREER-2239688 (D. W.), NCS-FO-2124179, and NCS-FO-1835231 (H. Pfister.); Office of Naval Research grant N00014-20-1-2828 (J. W. L). X. H. was supported by the Edward R. and Anne G. Lefler predoctoral fellowship from the Lefler Center for Neurodegenerative Disorders at Harvard Medical School and the Simmons Awards from the Harvard Center for Biological Imaging.

## Author Contribution

X. H., X. L., J. W. L, and J. S. T. conceived this study. J. S. T generated the mAb sequences. X. H. generated the scFv probes. T. F. and H. Ploegh provided materials for scFv probe production. X. H. performed the LM experiments. X. H. and R. S. performed the EM experiments. S. W. performed the imaging processing of the ssEM dataset. P. H. L. and V. J. performed 3D segmentation and curated the vCLEM dataset at Neuroglancer. Y. M. performed 2D segmentation. D. B. provided the reconstruction tool VAST and advice on 3D reconstruction and rendering. Y. W. helped with LM and EM co-registration. X. H., X. L., E. S. M., and S. A. performed 3D reconstruction and analysis. D. W., Z. L., J. A., and H. Pfister performed mitochondria and vesicle detection. X. H. and J. W. L wrote the paper with input from J. S. T., H. Ploegh., R. S., V. J., and P. H. L.

## Competing interests

The authors declare no competing interests.

## Notes

### Competing Interest Statement

The authors have declared no competing interest.

### Summary of Updates

Abstract and manuscript text edited to improve clarity; Corresponding authors updated

