## Supplemental Materials for "Multiplexed volumetric CLEM enabled by antibody derivatives provides new insights into the cytology of the mouse cerebellar cortex"

### Supplementary Materials

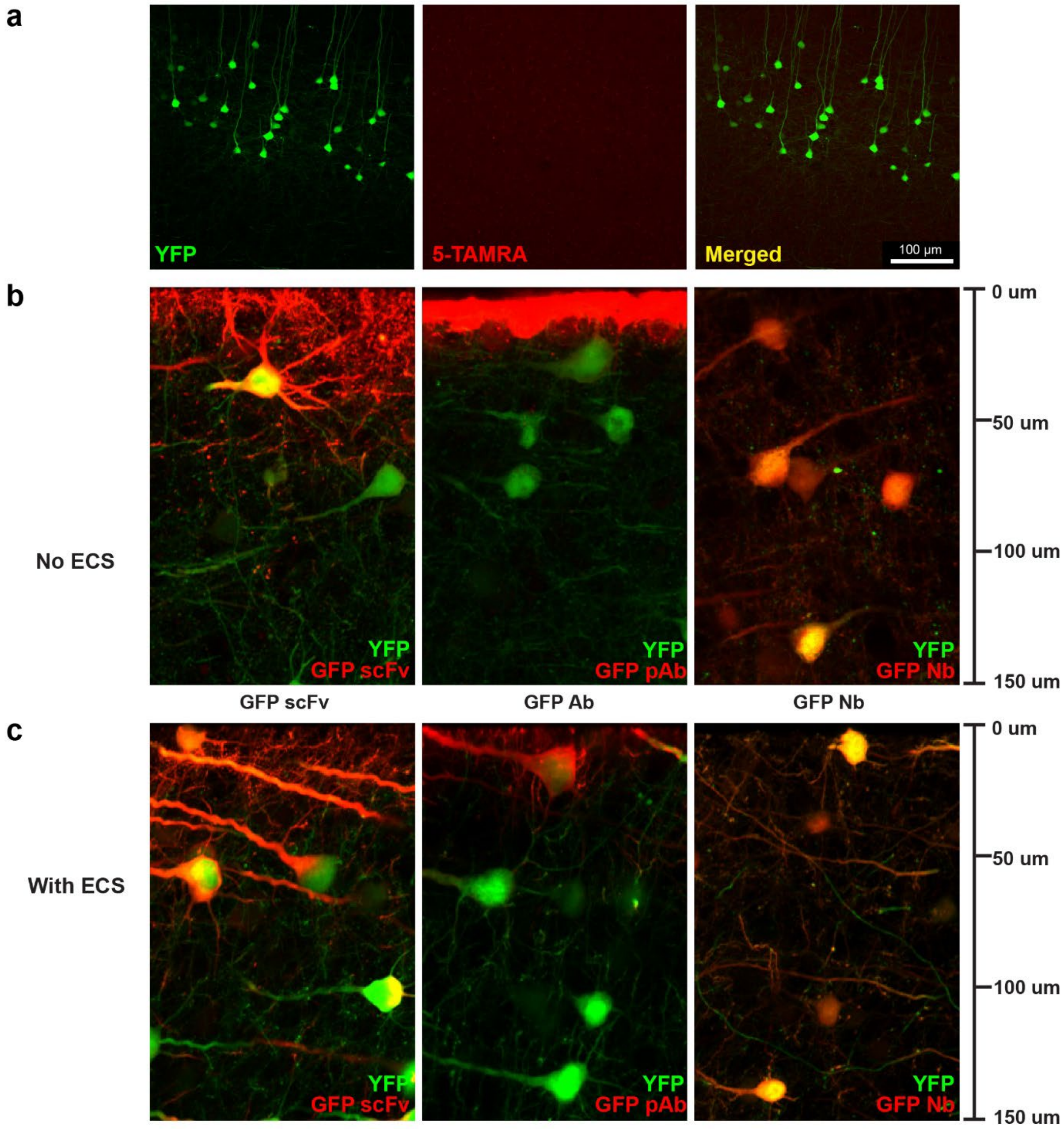


**Sup. Figure 1. Characterization of the GFP-specific scFv.**

**a**, Confocal images from the cerebral cortex of a unstained YFP-H mouse. **b** and **c**, Tissue penetration depth comparison of scFvs, pAbs, and nanobodies without or with the preservation of ECS.

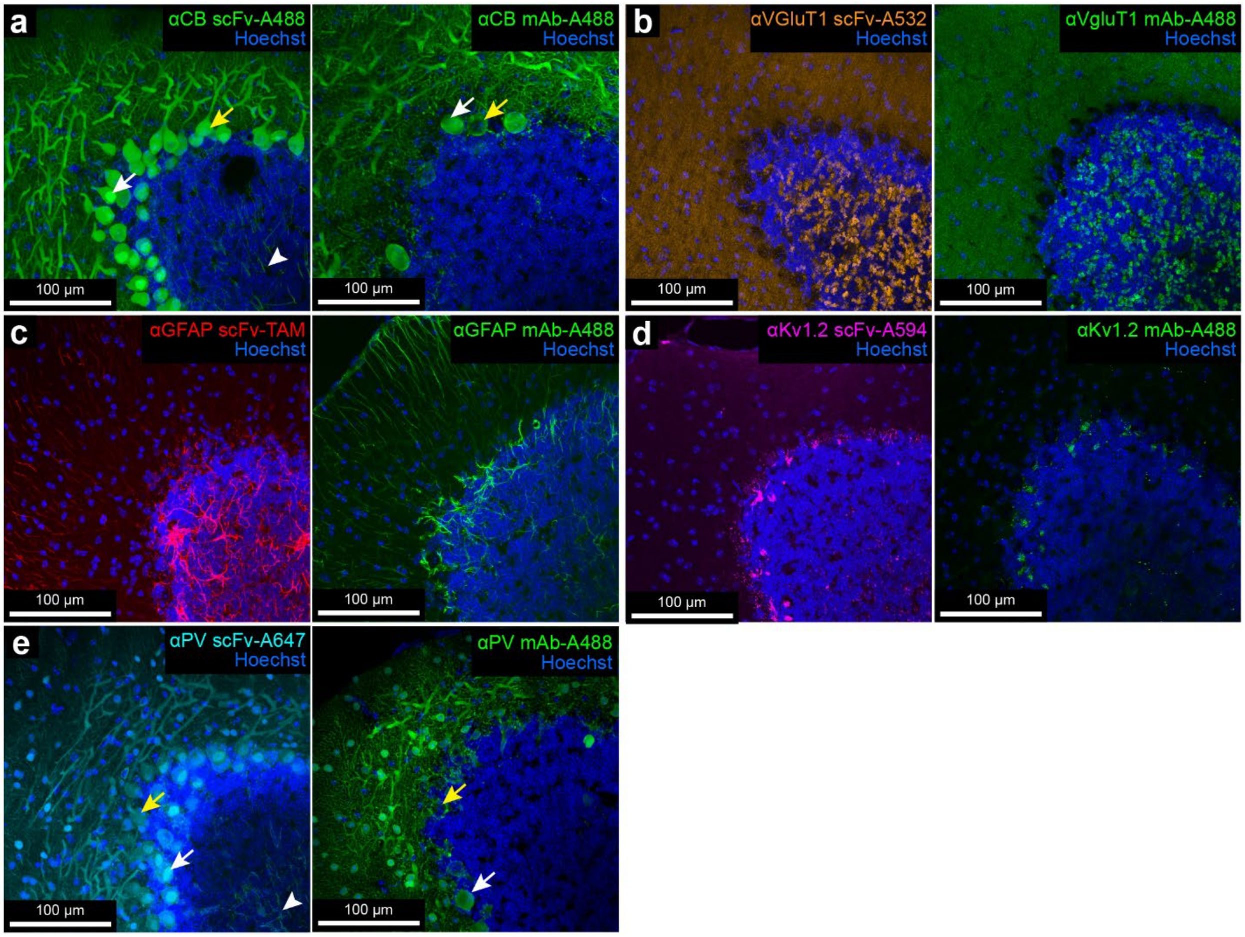


**Sup. Figure 2. Validation of immunofluorescence by scFv probes and their parental mAbs (part 1).**

**a**-**e**, Confocal images of different sections from the cerebellum Crus 1 stained with scFvs targeting CB, VGluT1, GFAP, Kv 1.2, and PV; or these five scFvs' parental mAbs and secondary antibodies conjugated with Alexa Fluor 488. White arrows in **a** and **e** indicate labeled cells that also had labeling in their nuclei. Yellow arrows in **a** and **e** indicate the labeled cell did not have labeling in their nuclei. White arrowheads in **a** and **e** indicate the labeled axons.

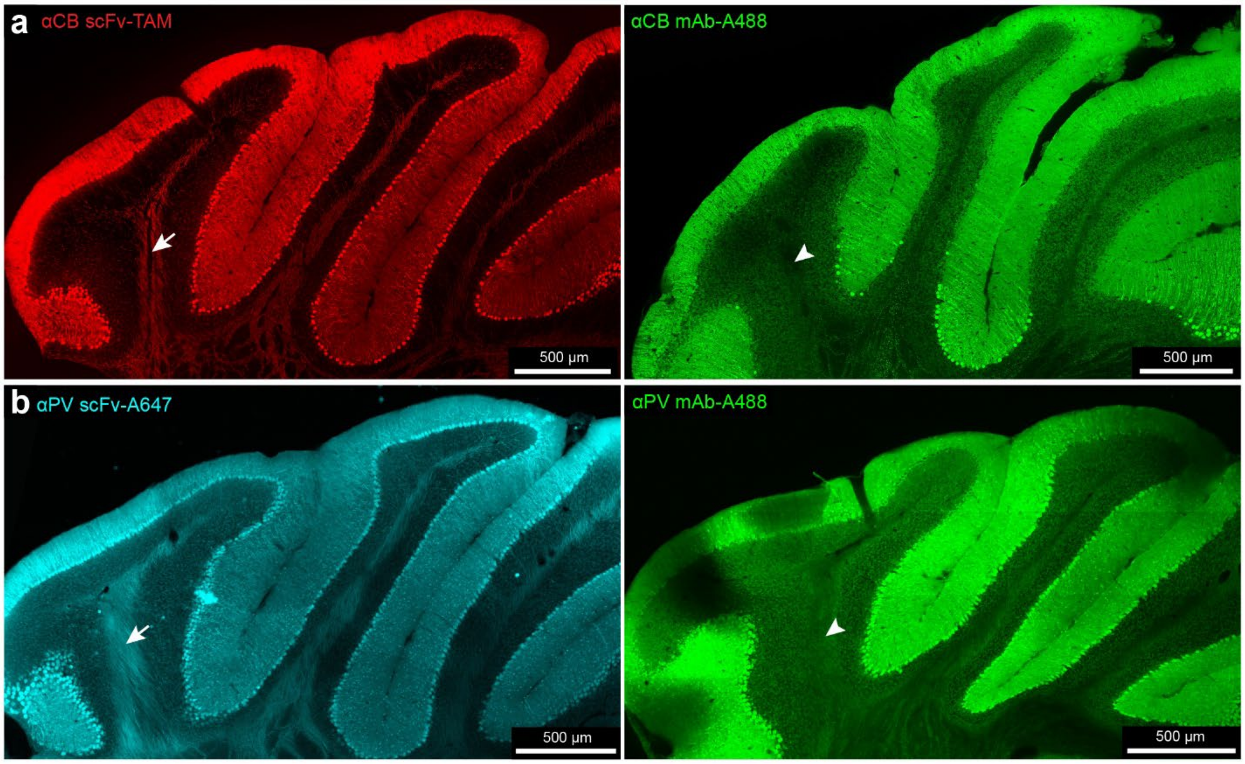


**Sup. Figure 3. Validation of immunofluorescence by CB-specific or PV-specific scFv probes.**

**a**-**b**, Confocal images of different coronal sections from the cerebellum stained with the CB-specific or the PV-sepcific scFv; or their parental mAbs and secondary antibodies conjugated with Alexa Fluor 488. White arrows in **a** and **b** indicate the labeled axons by the scFvs. White arrowheads in **a** and **e** indicate the unlabeled axons by the mAbs.

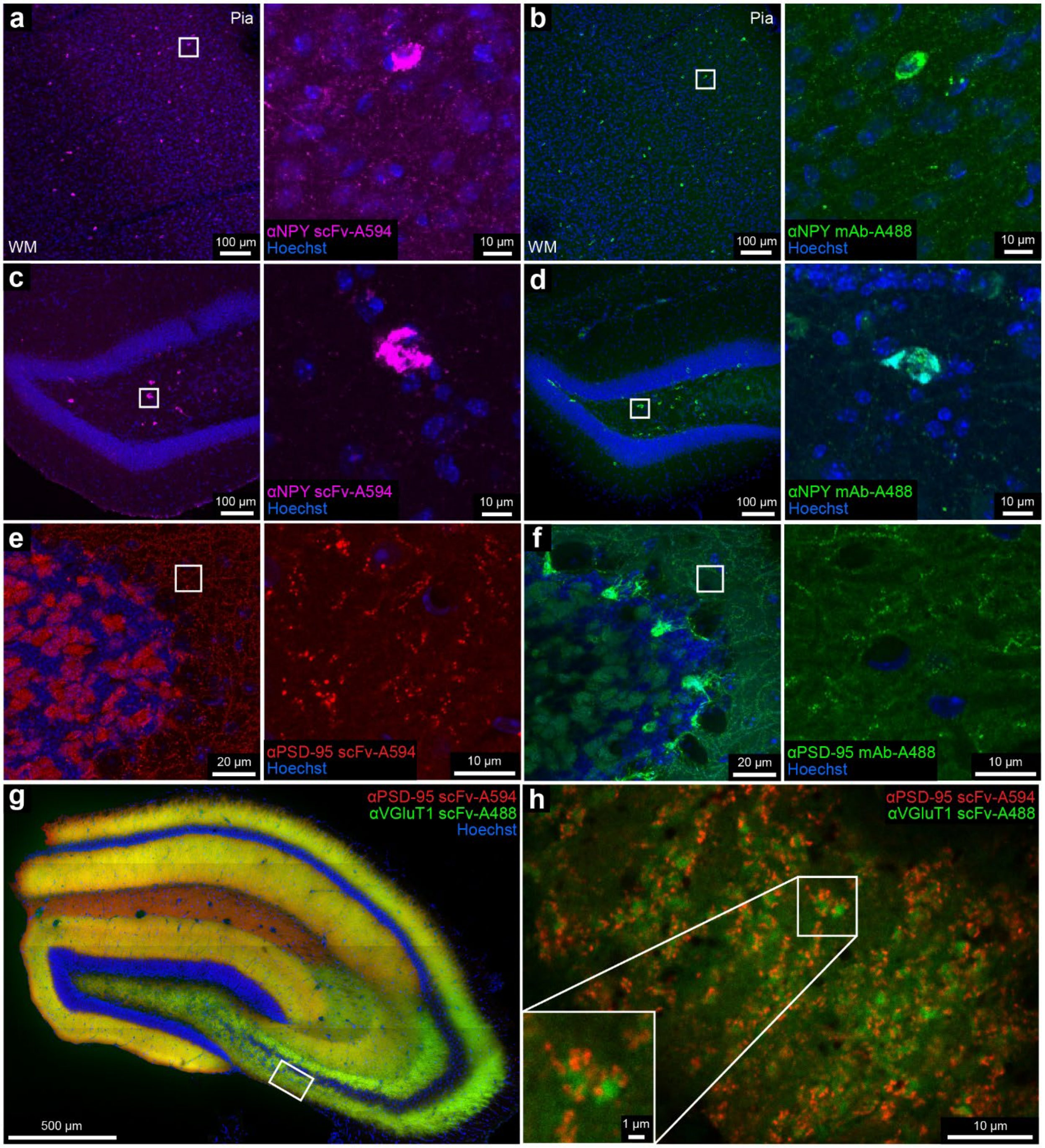


**Sup. Figure 4. Validation of immunofluorescence by NPY-specific or PSD-95-specific scFv probes.**

**a**-**f**, Confocal images of different sections from the cerebral cortex (**a** and **b**), the hippocampus (**c** and **d**), or the cerebellum Crus 1 (**e** and **f**) stained with scFvs targeting NPY or PSD-95; or their parental mAbs and secondary antibodies conjugated with Alexa Fluor 488. The right panels in **a**-**f** are enlarged boxed insets from the left panels. **g** and **h**, Confocal images of a section from hippocampus stained with scFvs targeting PSD-95 or VGluT1. **h** is the boxed inset from **g** imaged at higher magnification.

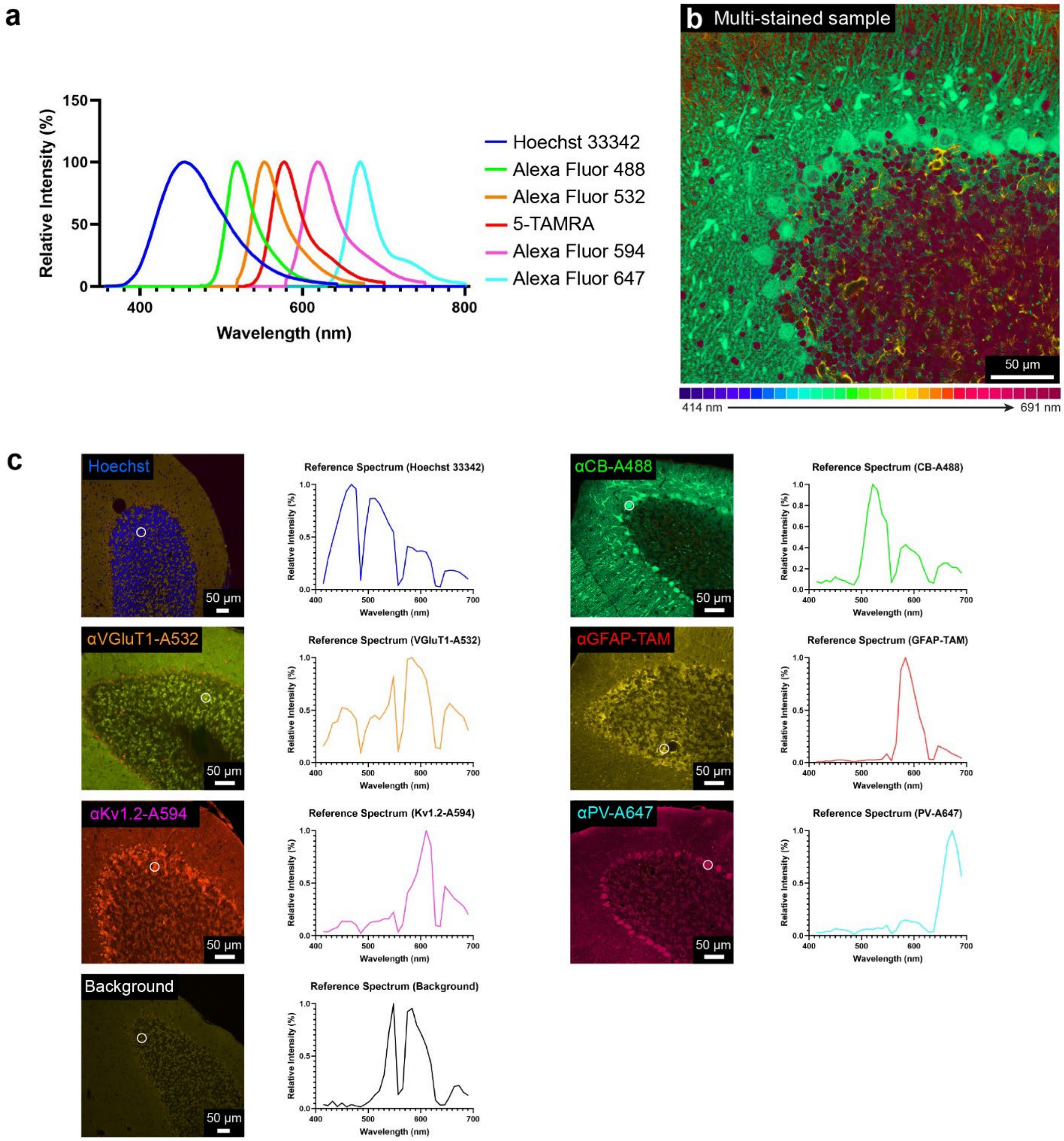


**Sup. Figure 5. Technical details of linear unmixing.**

**a**, The excitation and emission spectra of the six fluorescent dyes used in the multi-scFv labeling. **b**, An image slice from the lambda stack of the multicolor sample with a depth of 52 µm. The 32 channels (from 414 nm to 691 nm) in the lambda mode were labeled with different colors. **c**, the lambda stacks acquired from the individually labeled samples. The reference spectrum for each dye was extracted from the pixels labeled by the white circle in each lambda stack image.

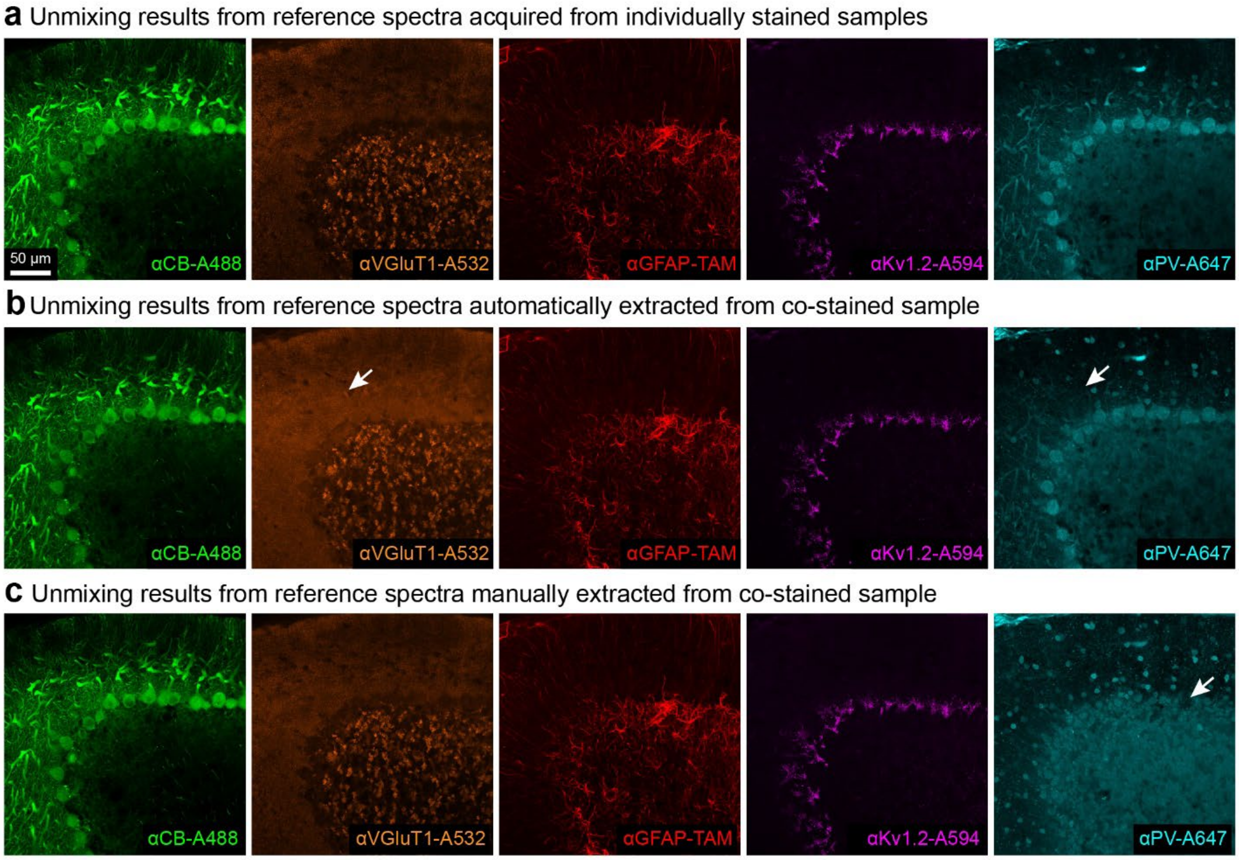


**Sup. Figure 6. Comparison of linear unmixing results using reference spectra extracted in three different ways.**

White arrows in **b** indicate the fluorescence signals that should be in the Alexa Fluor 647 channel were separated into the Alexa Fluor 532 channel instead. White arrows in **c** indicate where the fluorescence signals from the Purkinje cell bodies were missing in the Alexa Fluor 647 channel.

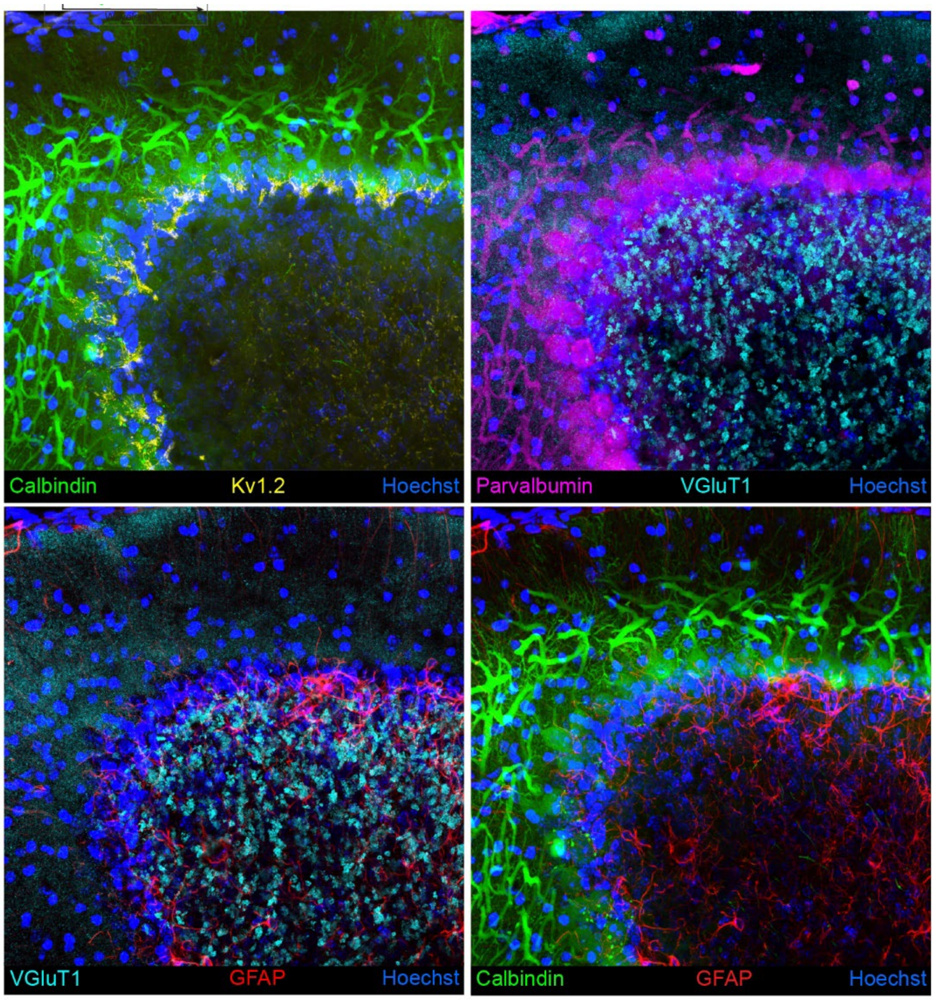


**Sup. Figure 7. Three-channel maximum intensity projection images of the multi-color fluorescence image stack in Figure 2 c.**

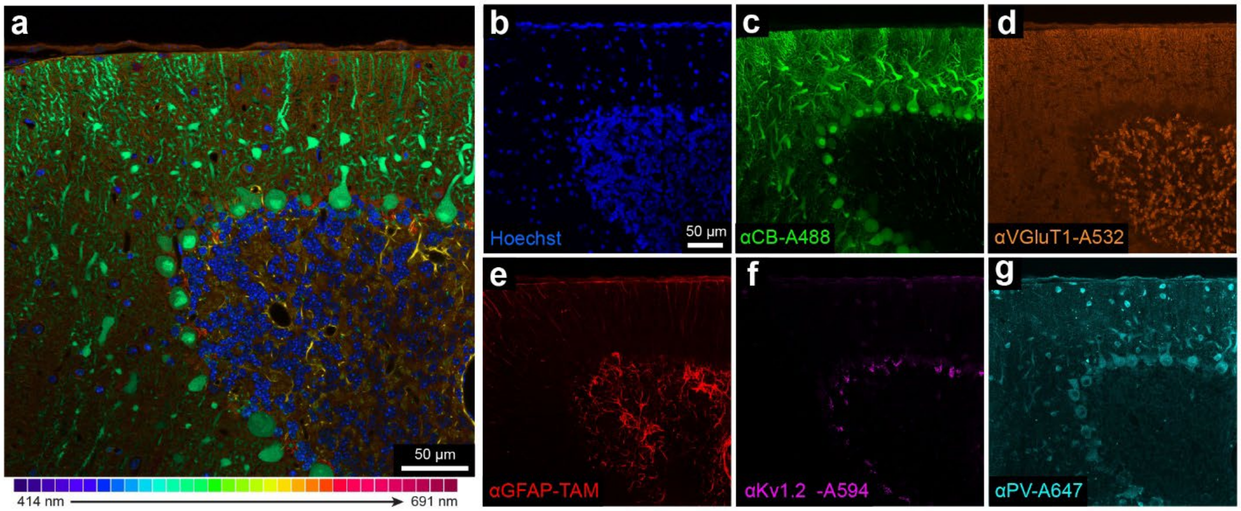


**Sup. Figure 8. The results of combining the Hoechst channel into linear unmixing of confocal micrographs.**

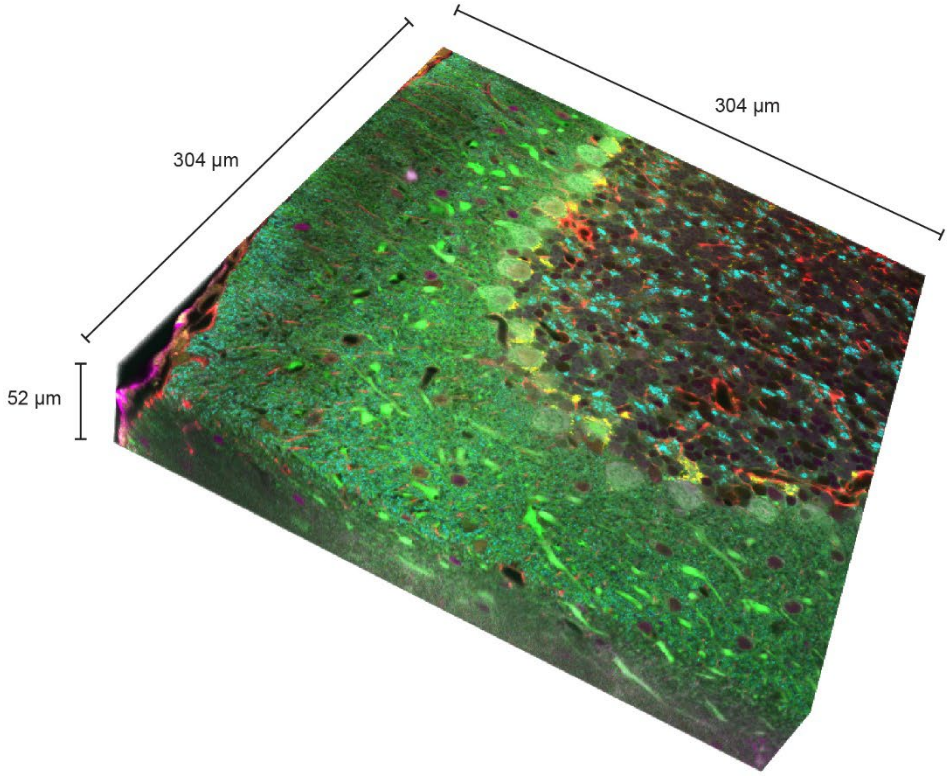


**Sup. Figure 9. The dimensions of the multi-color fluorescence image volume acquired by scFv-enabled immunofluorescence and linear unmixing.**

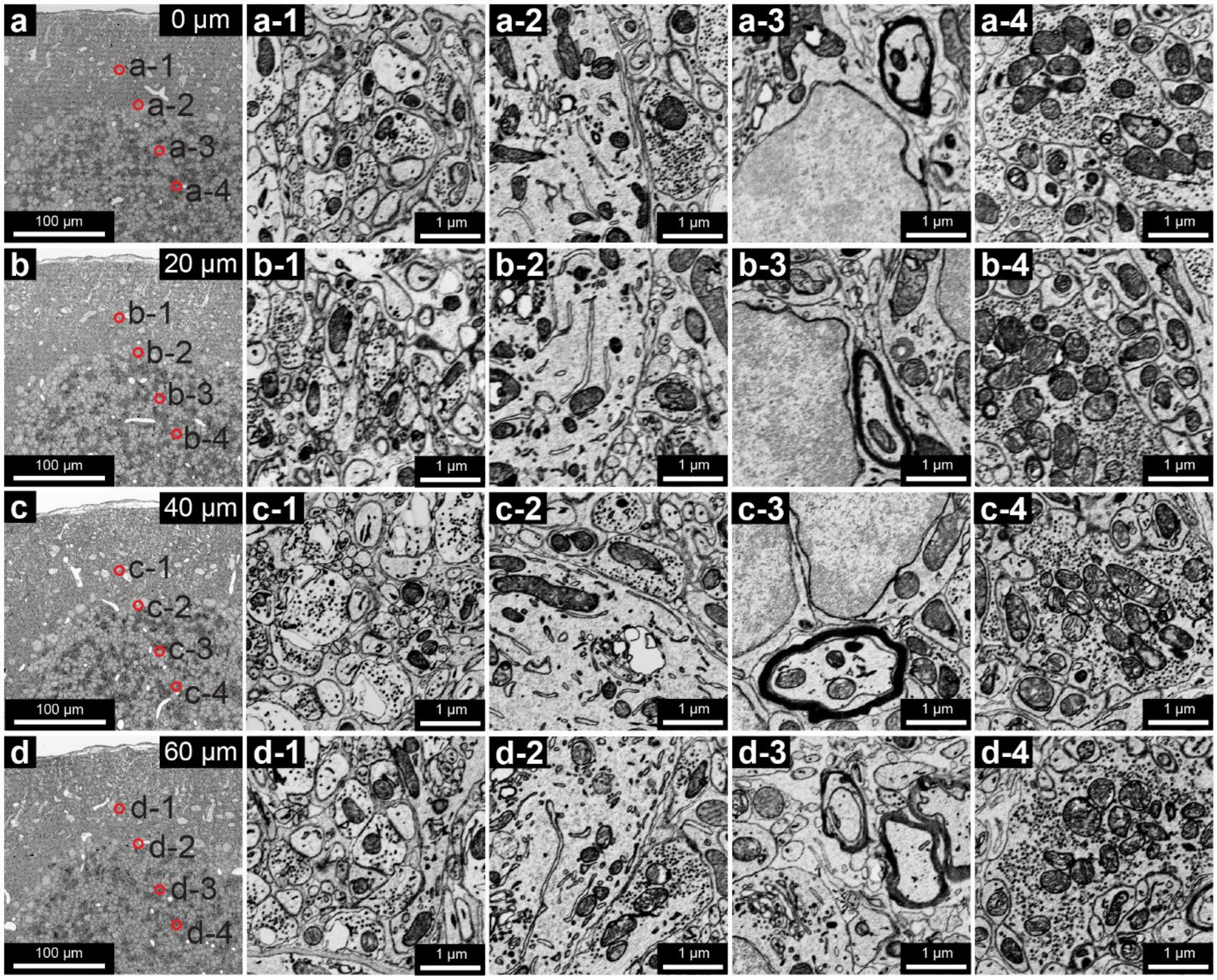


**Sup. Figure 10. Well-preserved ultrasctructure from the surface (a) to the middle (d) of the 120-µm section.**

Panel **1-4** in **a-d** show the ultrastructure at the locations labeled by the red circles in the right panels.

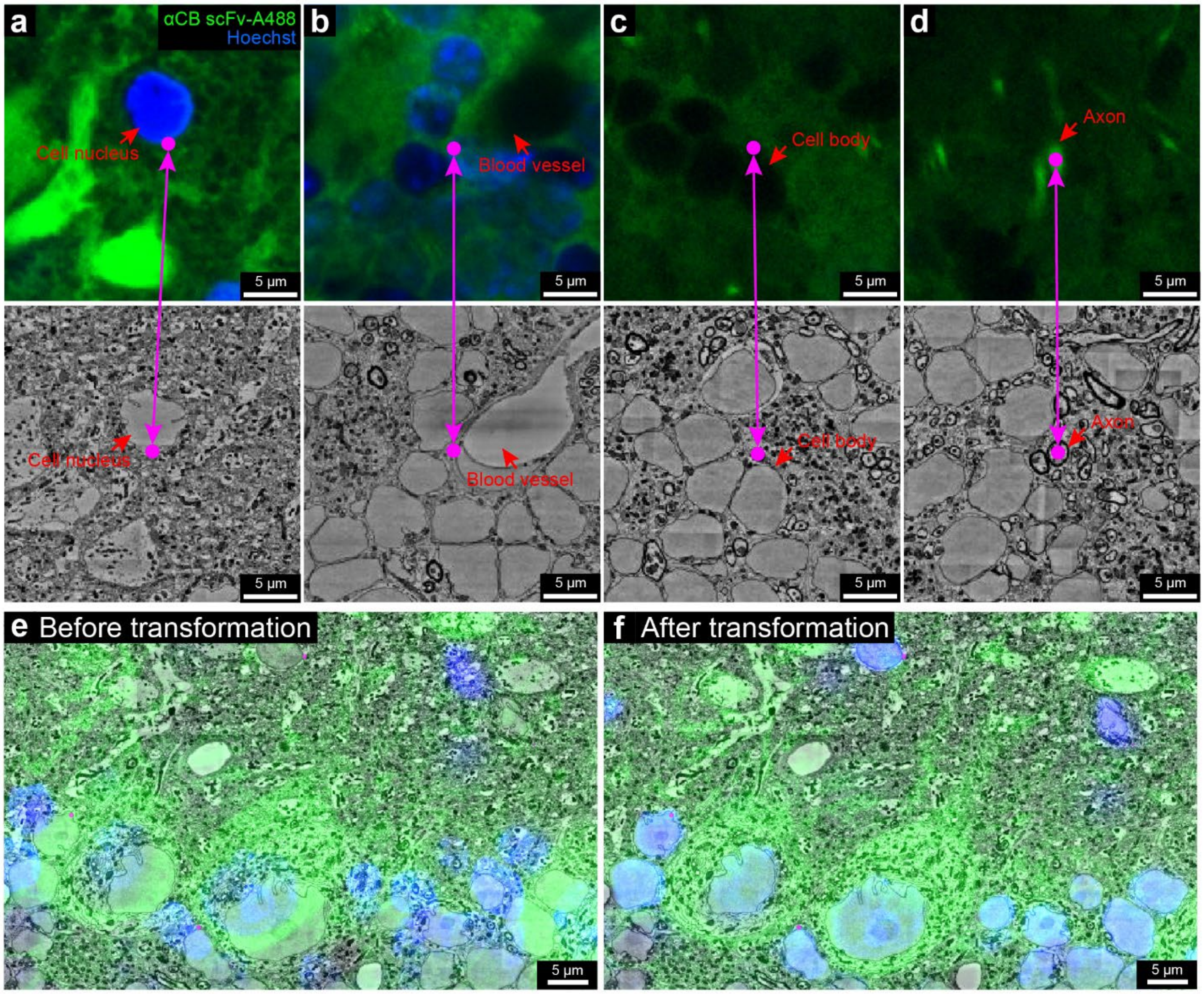


**Sup. Figure 11. Co-registration between fluorescence image volume and the high-resolution EM volume.**

**a**-**d,** landmark points that corresponded to the same sites in the two volumes were placed on blood vessels, cell nuclei, cell bodies, and axons. **e**-**f** show the overlays between the fluorescence image and the EM image before and after the transformation of the fluorescence image volume was performed based on the point correspondences.

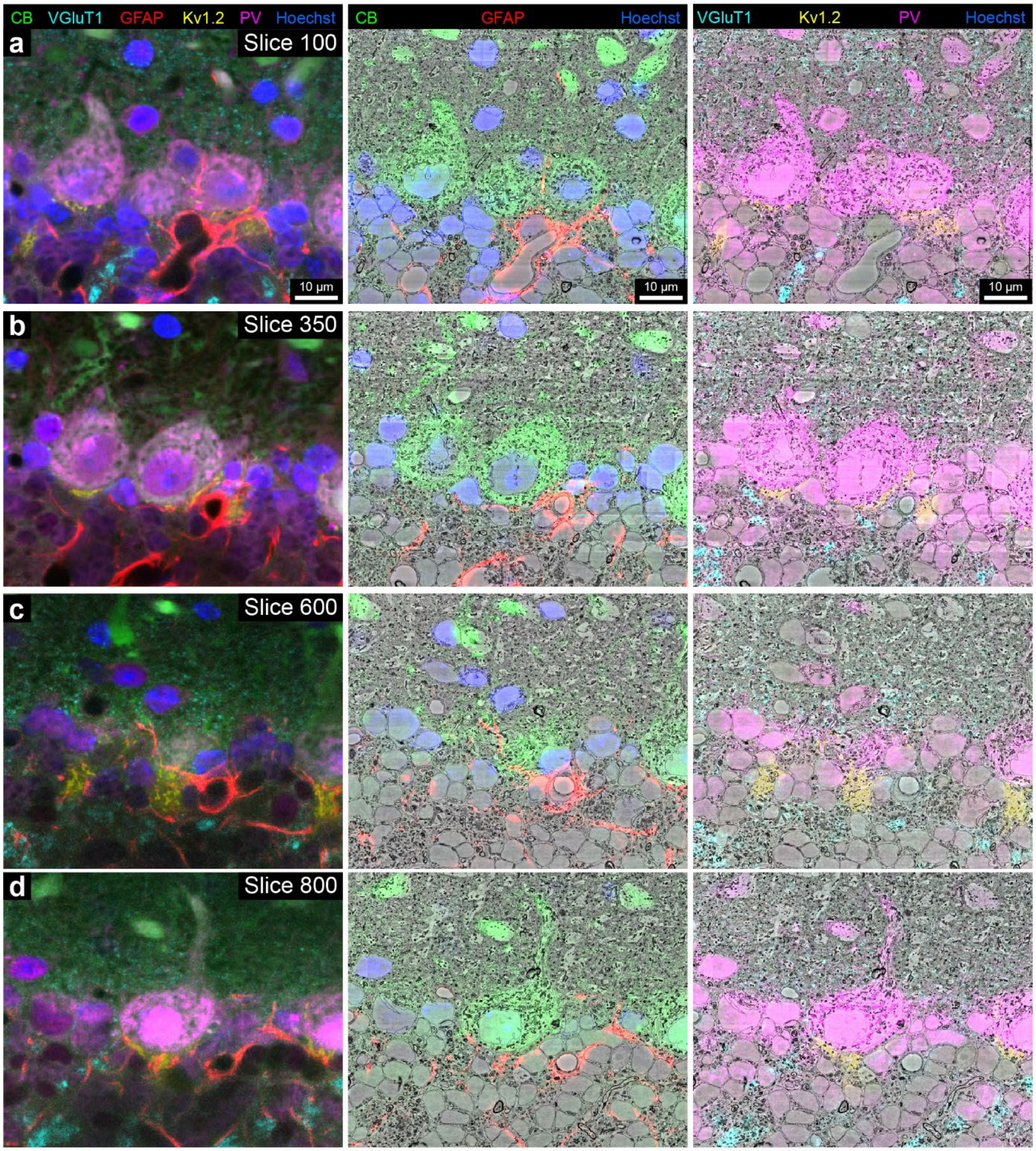


**Sup. Figure 12. Demonstration of the overlay between fluorescence signals and EM ultrastructure throughout the vCLEM dataset.**

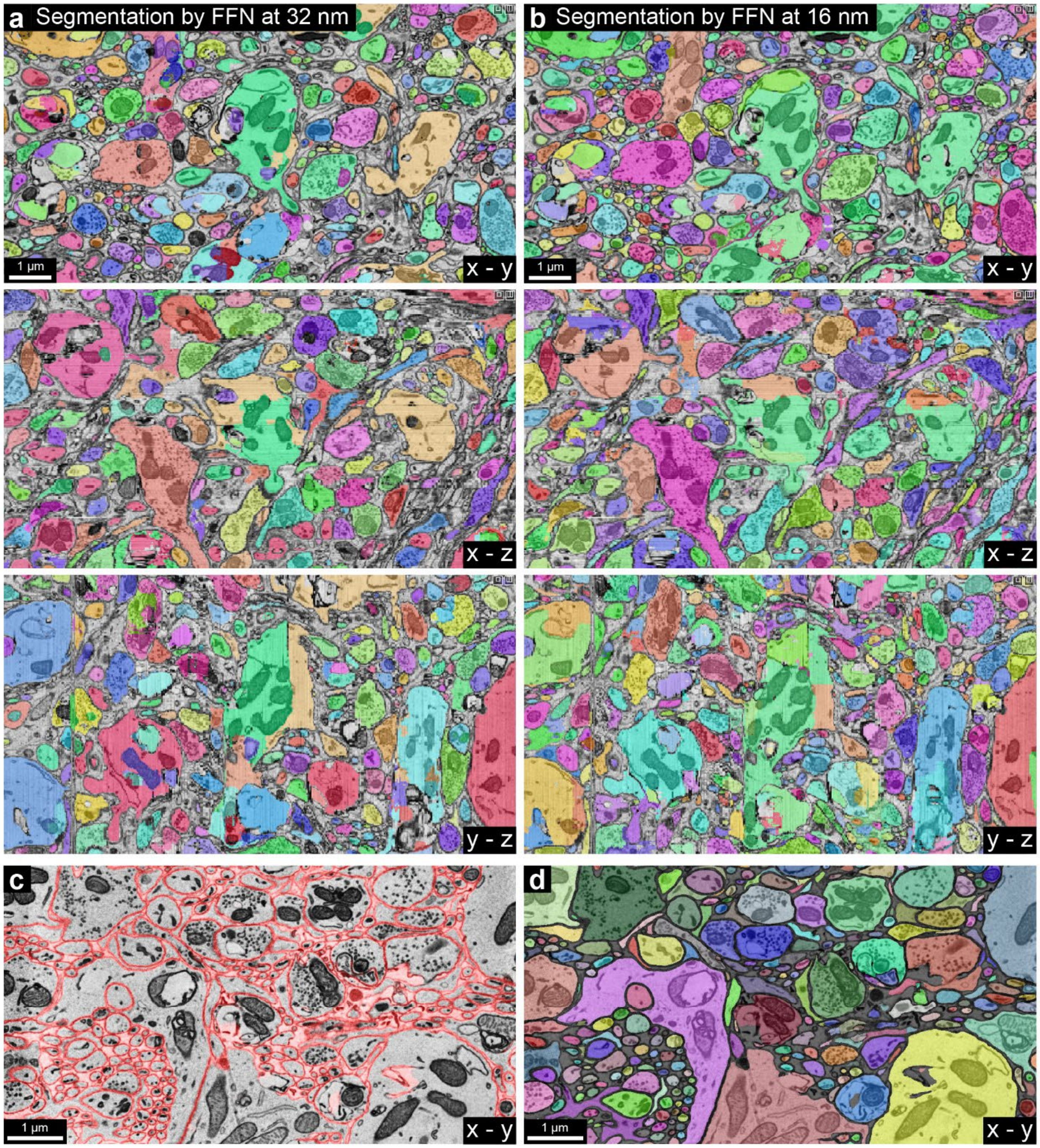


**Sup. Figure 13. Automatic segmentation results.**

3D segmentation results at 32 nm (**a**) and at 16 nm (**b**) by FFN [1]. Membrane prediction results at 8 nm (**c**) and 2D segmentation results at 8 nm (**d**) by a segmentation method developed by our lab [2].

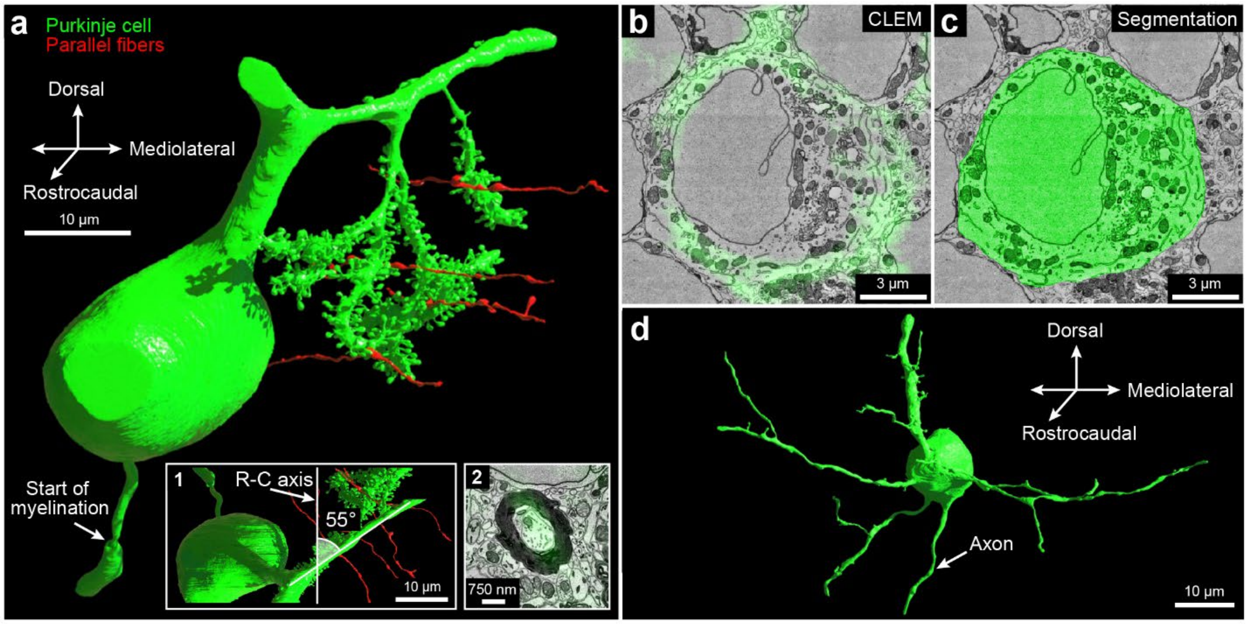


**Sup. Figure 14. 3D reconstruction of cells labeled by the calbindin-specific scFv probe.**

**a**, 3D reconstruction of the Purkinje cell labeled in Figure 4 a (green) and four parallel fibers (red) that made synapses on the Purkinje cell. Inset **1**, the reconstructed Purkinje cell viewed in the rostralcaudal-mediolateral plane. The dendritic tree of this Purkinje cell was not perpendicular to the rostral-caudal axis but intersected at the axis at an angle of around 55°. Inset **2**, the 2D CLEM image showing the fluorescence signal (green) of the calbindin-specific scFv probe overlaps with a heavily myelinated axon. **b**, the 2D CLEM image showing the fluorescence signal (green) of the calbindin-specific scFv probe overlaps with a Golgi cell. **c**, EM image showing the 2D segmentation (green) of the calbindin positive Golgi cell based on staining shown in **b**. **d**, 3D reconstruction of the Golgi cell labeled in **b**.

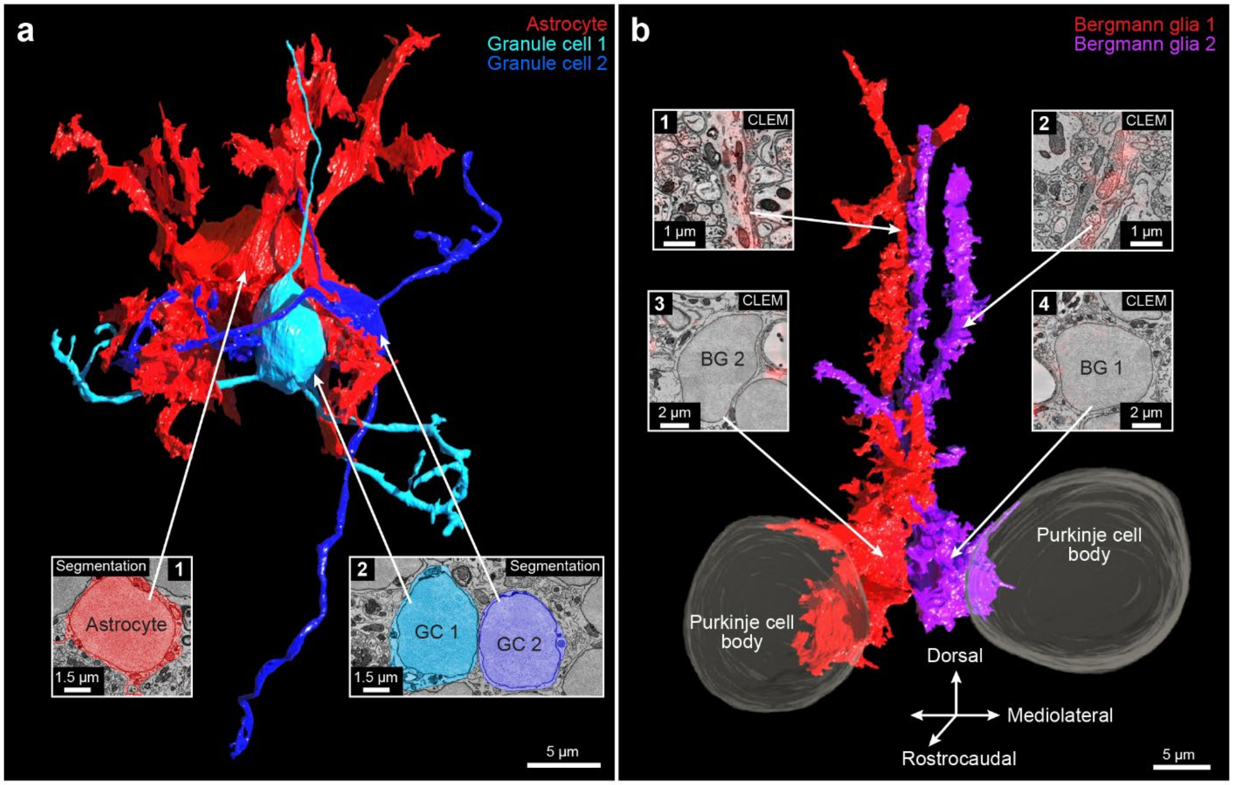


**Sup. Figure 15. 3D reconstruction of cells labeled by the GFAP-specific scFv probe.**

**a**, 3D reconstruction of the astrocyte labeled in Figure 4 e (red) and two nearby granule cells (cyan and blue). Insets, EM image showing the 2D segmentations of the cell bodies of the reconstructed astrocyte and granule cells. **b**, 3D reconstruction of the two labeled Bergmann glia labeled in Figure 4 i (red) and two nearby Purkinje cells. Insets **1** and **2**, the 2D CLEM images showing the fluorescence signal (red) of the GFAP-specific scFv probe overlaps with the processes of the Bergmann glia. Insets **3** and **4**, the 2D CLEM images showing the fluorescence signal (red) of the GFAP-specific scFv probe does not overlap with the cell bodies of the Bergmann glia.

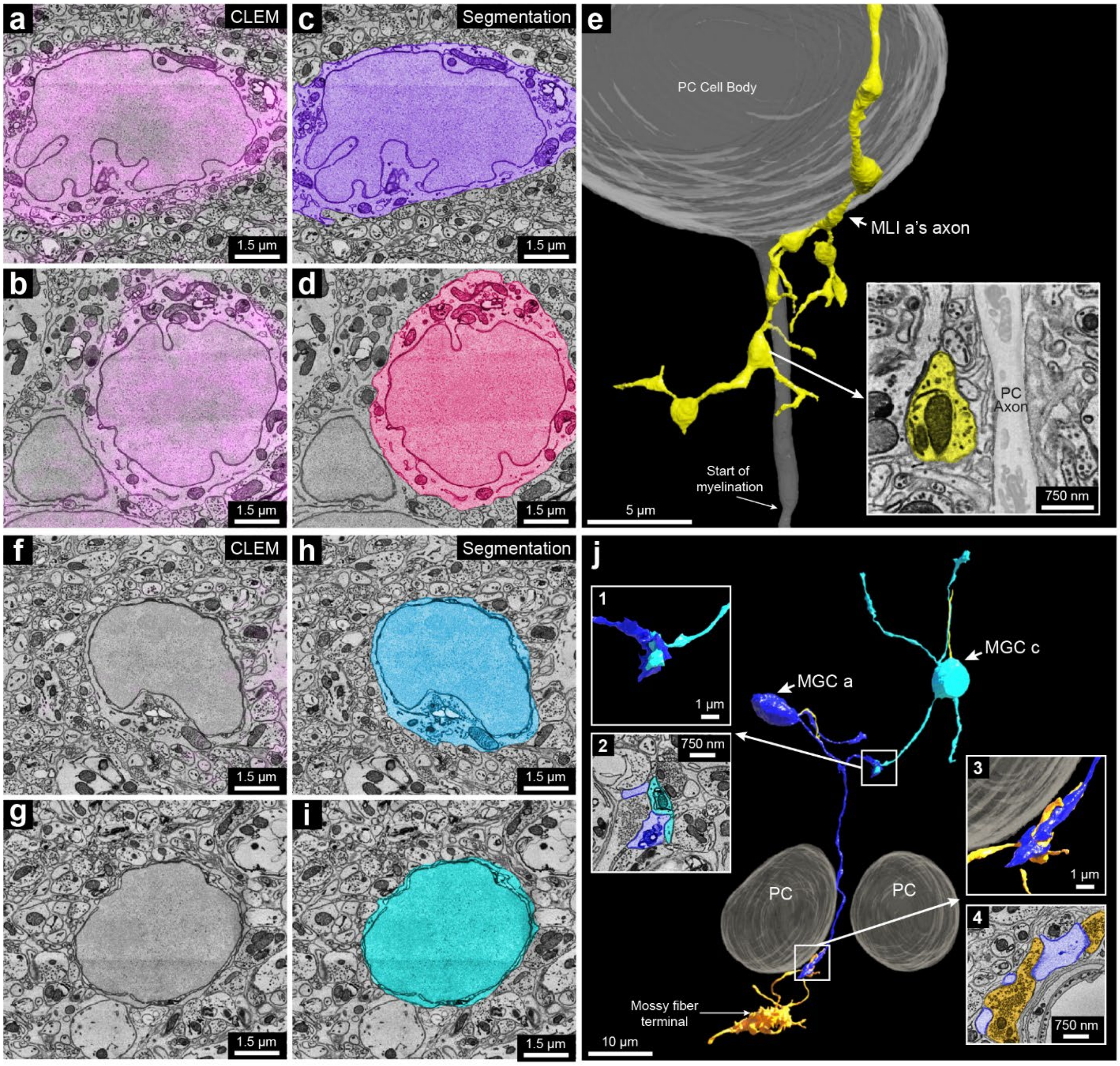


**Sup. Figure 16. 3D reconstruction of the molecular layer interneurons and granule cells.**

**a**-**b**, 2D CLEM image showing the fluorescence signals (magenta) of the parvalbumin-specific scFv probe overlap cell bodies of MLI b and c. **b**-**d**, EM image showing the 2D segmentation (magenta) of MLI a and b. **e**, the axon of MLI a was part of the pinceau structure that surrounds a Purkinje cell's axon initial segment (inset). **f**-**g**, 2D CLEM image showing the fluorescence signals (magenta) of the parvalbumin-specific scFv probe do not overlap cell bodies of MGC b and c. **h**-**i**, EM image showing the 2D segmentation (magenta) of MGC b and c. **j**, 3D reconstruction MGC a and MGC c. These two cells formed a glomerulus (insets **1** and **2**). MGC a received synaptic inputs from a mossy fiber terminal (insets **3** and **4**).

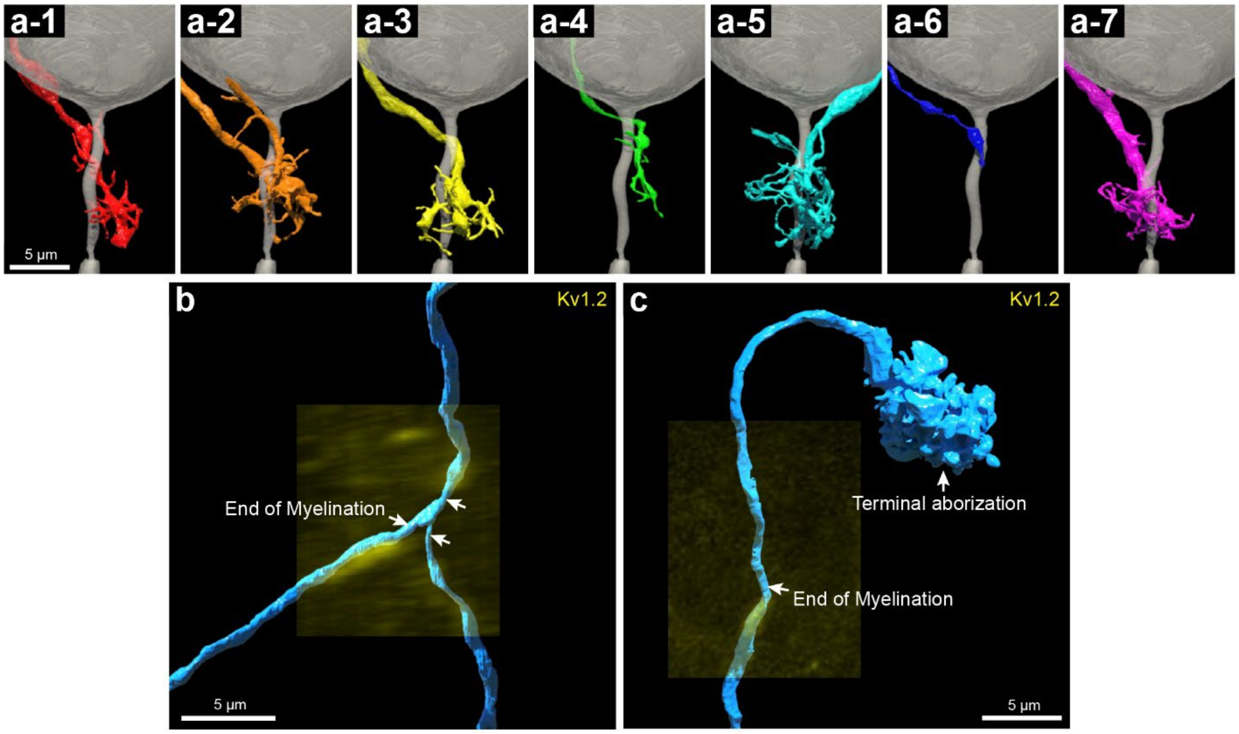


**Sup. Figure 17. 3D reconstruction of axonal terminals in a pinceau and the mossy fibers labeled with the Kv 1.2-specific scFv.**

**a-1** to **a-7**, seven individual axon terminals labeled by the Kv 1.2-specific scFv probe at the pinceau structure in Figure 6 a. **b**, 3D reconstruction of an axon with Kv1.2 positive juxtaparanodal labeling (yellow fluorescence) at a branching point. **c**, 3D reconstruction of an axon with Kv1.2 positive juxtaparanodal labeling (yellow fluorescence) at the site where the myelination ended. This axon ended in a terminal axonal arborization (arrow) in the granule cell layer.

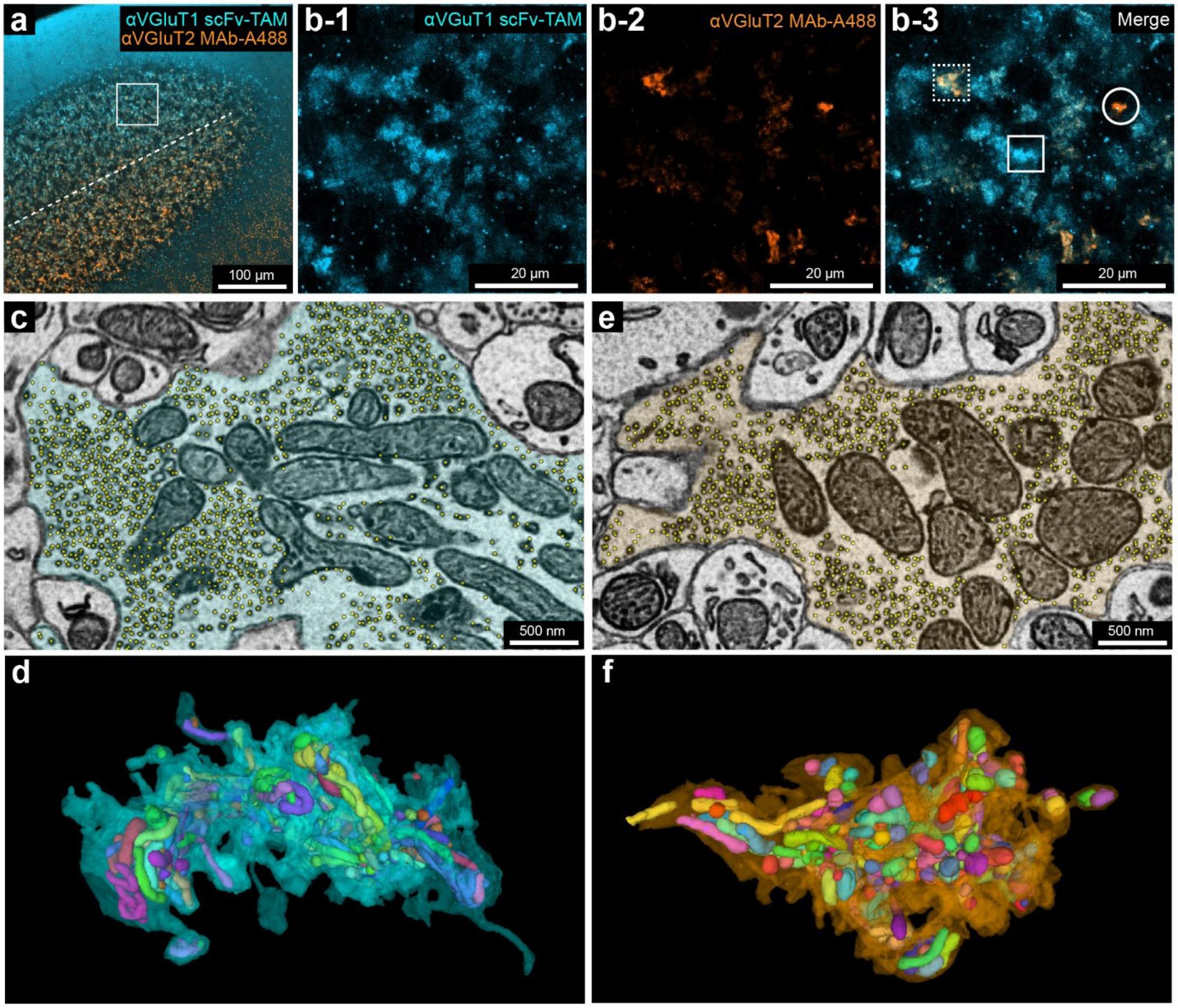


**Sup. Figure 18. Three types of mossy fiber terminals identified by double immunofluorescence of VGluT1/2 and synaptic vesicle and mitochondria detection results.**

**a**, Confocal image of a section from the cerebellum Crus 1 stained with a VGluT1-specific scFv probe conjugated with Alexa Fluor 5-TAMRA and a VGluT2-specific scFv probe with secondary antibodies conjugated with Alexa Fluor 488. The image was pseudo-colored. Note the labeling of VGLUT1 was stronger than labeling of VGluT2 at the left side of the dotted line and vice versa. **b-1** to **b-3**, Single-channel and two-channel images enlarged from the boxed inset in **a**. The box with solid lines indicates a VGluT1 positive terminal. The box with dotted lines indicates a VGluT1/2 double positive terminal. The circle indicates a VGluT2 positive terminal. **c**-**d,** The synaptic vesicle and mitochondria detection results from a reconstructed VGluT1 positive terminal. Colored objects inside the transparent terminal are the detected mitochondria. **e**-**f,** the synaptic vesicle and mitochondria detection results from a reconstructed VGluT2 positive terminal. Colored objects inside the transparent terminal are the detected mitochondria.

**Sup. Table 1. The final concentrations of scFv and nanobody probes used in the immunofluorescence of this work.**

| Target | Clone No. | Fluorescent dyes conjugated | Final conc. |
|---|---|---|---|
| **GFP (scFv)** | N86/38 | 5-TAMRA | 0.01 mg/ml |
| **GFP (nanobody)** | Enh [3] | Alexa Fluor 647 | 0.01 mg/ml |
| **Calbindin** | L109/57 | 5-TAMRA, Alexa Fluor 488 | 0.01 mg/ml |
| **GFAP** | N206B/9 | 5-TAMRA | 0.01 mg/ml |
| **VGluT1** | N28/9 | 5-TAMRA, Alexa Fluor 532 | 0.015 mg/ml |
| **PSD-95** | K28/43 | 5-TAMRA | 0.01 mg/ml |
| **Kv1.2** | K14/16 | Alexa Fluor 594 | 0.02 mg/ml |
| **Parvalbumin** | L114/81 | Alexa Fluor 647 | 0.01 mg/ml |
| **Neuropeptide Y** | L115/13 | Alexa Fluor 647 | 0.01 mg/ml |

**Sup. Table 2.The dilution ratios of primary antibodies used in the immunofluorescence of used in this work.**

| Target | Clone No. | Vendor | Catalog No. | Dilution Ratio |
|---|---|---|---|---|
| **GFP** | N.A. | ThermoFisher | A-31852 | 1:200 |
| **Calbindin** | L109/57 | Antibodies Incorporated | 75-448 | 1:200 |
| **GFAP** | N206B/9 | Antibodies Incorporated | 75-279 | 1:200 |
| **VGluT1** | N28/9 | Antibodies Incorporated | 75-066 | 1:200 |
| **VGluT2** | N28/29 | Antibodies Incorporated | 75-067 | 1:200 |
| **PSD-95** | K28/43 | Antibodies Incorporated | 75-028 | 1:200 |
| **Kv1.2** | K14/16 | Antibodies Incorporated | 75-008 | 1:500 |
| **Parvalbumin** | L114/81 | Antibodies Incorporated | 75-479 | 1:200 |
| **Neuropeptide Y** | L115/13 | Antibodies Incorporated | 75-456 | 1:200 |

**Sup. Table 3. The final concentrations of secondary antibodies used in the immunofluorescence of used in this work.**

| Name | Fluorescence dye conjugated | Vendor | Catalog No. | Dilution Ratio |
|---|---|---|---|---|
| **Goat anti-Mouse IgG1 Secondary Antibody** | Alexa Fluor 488 | ThermoFisher | A-21121 | 1:100 |
| **Goat anti-Mouse IgG2a Secondary Antibody** | Alexa Fluor 488 | ThermoFisher | A-21131 | 1:100 |
| **Goat anti-Mouse IgG2b Secondary Antibody** | Alexa Fluor 488 | ThermoFisher | A-21141 | 1:100 |

**Sup. Table 4. Information on the VGluT1 positive terminals.**

| Positive terminal No. | Volume / $\mu m^3$ | Vesicle Number | Vesicle density per $\mu m^3$ | Mito volume / $\mu m^3$ | Mito volume ratio |
|---|---|---|---|---|---|
| **1** | 123.19 | 436042 | 3540 | 24.13 | 19.59% |
| **2** | 109.43 | 426878 | 3901 | 22.04 | 20.14% |
| **3** | 85.79 | 292704 | 3412 | 20.81 | 24.26% |
| **4** | 124.89 | 409789 | 3281 | 30.94 | 24.77% |
| **5** | 97.90 | 233785 | 2388 | 29.45 | 30.08% |
| **6** | 89.89 | 255883 | 2847 | 22.09 | 24.57% |
| **7** | 92.86 | 277102 | 2984 | 22.19 | 23.90% |
| **8** | 84.47 | 246042 | 2913 | 24.43 | 28.92% |
| **9** | 75.39 | 269515 | 3575 | 15.90 | 21.09% |
| **10** | 109.52 | 293002 | 2675 | 31.37 | 28.64% |

**Sup. Table 5. Information on the VGluT1 negative terminals.**

| Negative terminal No. | Volume / µm³ | Vesicle Number | Vesicle density per µm³ | Mito volume / µm³ | Mito volume ratio |
|---|---|---|---|---|---|
| **1** | 71.81 | 196695 | 2739 | 20.56 | 28.63% |
| **2** | 92.23 | 268178 | 2812 | 24.62 | 26.70% |
| **3** | 75.40 | 144275 | 1913 | 24.54 | 32.55% |
| **4** | 33.73 | 97526 | 2891 | 10.56 | 31.31% |
| **5** | 151.10 | 424845 | 2812 | 43.92 | 29.07% |
| **6** | 61.82 | 142368 | 2303 | 17.92 | 29.00% |
| **7** | 40.06 | 116529 | 2909 | 9.03 | 22.54% |
| **8** | 68.55 | 213897 | 3120 | 16.70 | 24.36% |
| **9** | 40.73 | 120864 | 2968 | 10.50 | 25.78% |
| **10** | 42.79 | 141617 | 3309 | 9.37 | 21.90% |
